# A CHCHD10 variant causing ALS elicits an unfolded protein response through the IRE1/XBP1 pathway

**DOI:** 10.1101/2020.05.05.078881

**Authors:** Isabella R. Straub, Woranontee Weraarpachai, Eric A. Shoubridge

## Abstract

Mutations in *CHCHD10*, coding for a mitochondrial intermembrane space protein, are a rare cause of autosomal dominant amyotrophic lateral sclerosis (ALS). Mutation-specific toxic gain of function or haploinsuffuciency models have been proposed to explain pathogenicity. To decipher the metabolic dysfunction associated with the haploinsufficient p.R15L variant we integrated transcriptomic, metabolomic and proteomic data sets in patient cells subjected to nutrient stress. Patient cells had a complex I deficiency resulting in an increased NADH/NAD^+^ ratio, downregulation of the TCA cycle, and a reorganization of one carbon metabolism. This led to phosphorylation of AMPK, activation of an endoplasmic reticulum and mitochondrial unfolded protein response (UPR), and the production of GDF15 and FGF21, which are markers of mitochondrial disease. The endoplasmic reticulum UPR was mediated through the IRE1/XBP1 pathway, and was accompanied by reduced eIF2alpha phosphorylation, dephosphorylation of both JNK isoforms, and up regulation of several dual specific phosphatases. This study demonstrates that loss of CHCHD10 function elicits a striking energy deficit that activates cellular stress pathways, which may underlie the selective vulnerability of motor neurons.

## Introduction

Autosomal dominant mutations in coiled-helix coiled-helix domain containing protein 10 (*CHCHD10*) were recently identified as rare genetic causes of ALS (Bannwarth, Ait-El-Mkadem et al., 2014, Chaussenot, Le Ber et al., 2014, Chio, Mora et al., 2015, Dols-Icardo, Nebot et al., 2015, Johnson, Glynn et al., 2014, Kurzwelly, Kruger et al., 2015, Lehmer, Schludi et al., 2018, Muller, Andersen et al., 2014, Ronchi, Riboldi et al., 2015, Ryan, Zaldivar Vaillant et al., 2019, Shen, He et al., 2017, Zhang, Xi et al., 2015, Zhou, Chen et al., 2017). Most predicted pathogenic variants are present in the N-terminal half of the protein and are rarely found in the defining CHCH-domain in the C-terminus. The mutation c.44C>A (predicting p. R15L) has been reported in sporadic and familial ALS, and motor neuron disease in four studies (Johnson et al., 2014, Khan, Nikkanen et al., 2017, Muller et al., 2014, Zhang et al., 2015). CHCHD10 is a soluble 14 kDa mitochondrial protein which is upregulated in stress conditions, and localizes to the mitochondrial intermembrane space where it forms a complex with its paralogue CHCHD2 (Huang, Wu et al., 2018, Straub, Janer et al., 2018). However, the precise function of CHCHD10 remains unknown.

Mitochondrial dysfunction has long been suggested to contribute to ALS disease pathology, but prior to the identification of mutations in CHCHD10, these deficiencies were thought to be pleiotropic effects. Although all reported pathogenic mutations in CHCHD10 are autosomal dominant, the molecular basis of pathogenicity of CHCHD10 is variable. For instance, the p.S59L variant reported in patients with ALS-FTD (frontotemporal dementia) is associated with a fragmented mitochondrial network, protein aggregates in mitochondria, and the activation of the integrated stress response (ISR), ascribed to a toxic gain of function (Anderson, Bredvik et al., 2019, Bannwarth et al., 2014, Genin, Bannwarth et al., 2018, Genin, Madji Hounoum et al., 2019). On the other hand, the p.R15L and p.G66V variants, in which reduced levels of CHCHD10 protein are associated with mitochondrial respiratory chain dysfunction, appear to be haploinsufficient (Brockmann, Freischmidt et al., 2018, Penttila, Jokela et al., 2015, Straub et al., 2018).

Emerging lines of evidence suggest a link between the endoplasmic reticulum (ER) unfolded protein response and neurodegenerative diseases like ALS (Hetz & Mollereau, 2014). This unfolded protein stress response is designed to maintain and recover ER proteostasis through three distinct signalling pathways: PRKR-like endoplasmic reticulum kinase (PERK), activating transcription factor 6 (ATF6), and inositol-requiring kinase 1 (IRE1) (Hughes & Mallucci, 2019). In particular the IRE1 signalling cascade has been suggested as potential target for treatment as downstream targets such as XBP1 and CHOP have been shown to play a role in several ALS disease models, and knockdown of XBP1 provides significant protection against neuronal death and delays onset of disease (Hetz, Thielen et al., 2009, Ito, Yamada et al., 2009, Jaronen, Goldsteins et al., 2014, Medinas, Gonzalez et al., 2017).

To investigate the cellular and metabolic remodelling caused by the CHCHD10 variant p.R15L, we collected metabolomic, transcriptomic and proteomic datasets on patient fibroblasts. We cultured patient and rescued cells (expressing wild-type CHCHD10) in medium containing either glucose or galactose as a nutrient source. Glucose-free galactose medium imposes a nutrient stress, as it forces cells to rely solely on oxidative phosphorylation (OXPHOS) for energy production. Our data demonstrate a global remodelling of mitochondrial and cellular metabolic pathways, and the activation of an unfolded protein (UPR) stress response mediated by the IRE1/XBP1 pathway in the ER, and by ATF4 and ATF5 in mitochondria, in patient cells under nutrient stress conditions.

## Results

To uncover the mechanisms of pathogenesis in patient fibroblasts with the CHCHD10 p.R15L variant (hereafter referred to as ‘patient’), we set to compare them to patient cells expressing wild-type CHCHD10 cDNA (hereafter called ‘rescue’). We previously showed that about 20% of patient cells died over 24 hours after transfer to galactose medium, while the remaining cells remained attached to the plate, but did not re-enter the cell cycle. This phenotype could be rescued by expression of wild-type CHCHD10 cDNA. To discover the changes in patient cells, the transcriptome, proteome, and metabolome of patient and rescue cells were determined by RNA sequencing, tandem mass tag (TMT) mass spectrometry, and targeted metabolomics analysis in response to the galactose challenge. We compared the patient to the rescued cells grown in glucose medium (‘Glucose’) or in galactose medium for 48 hours (‘Galactose’) to profile the initial response to the galactose challenge in the patient (Fig. 1A).

**Figure 1.**
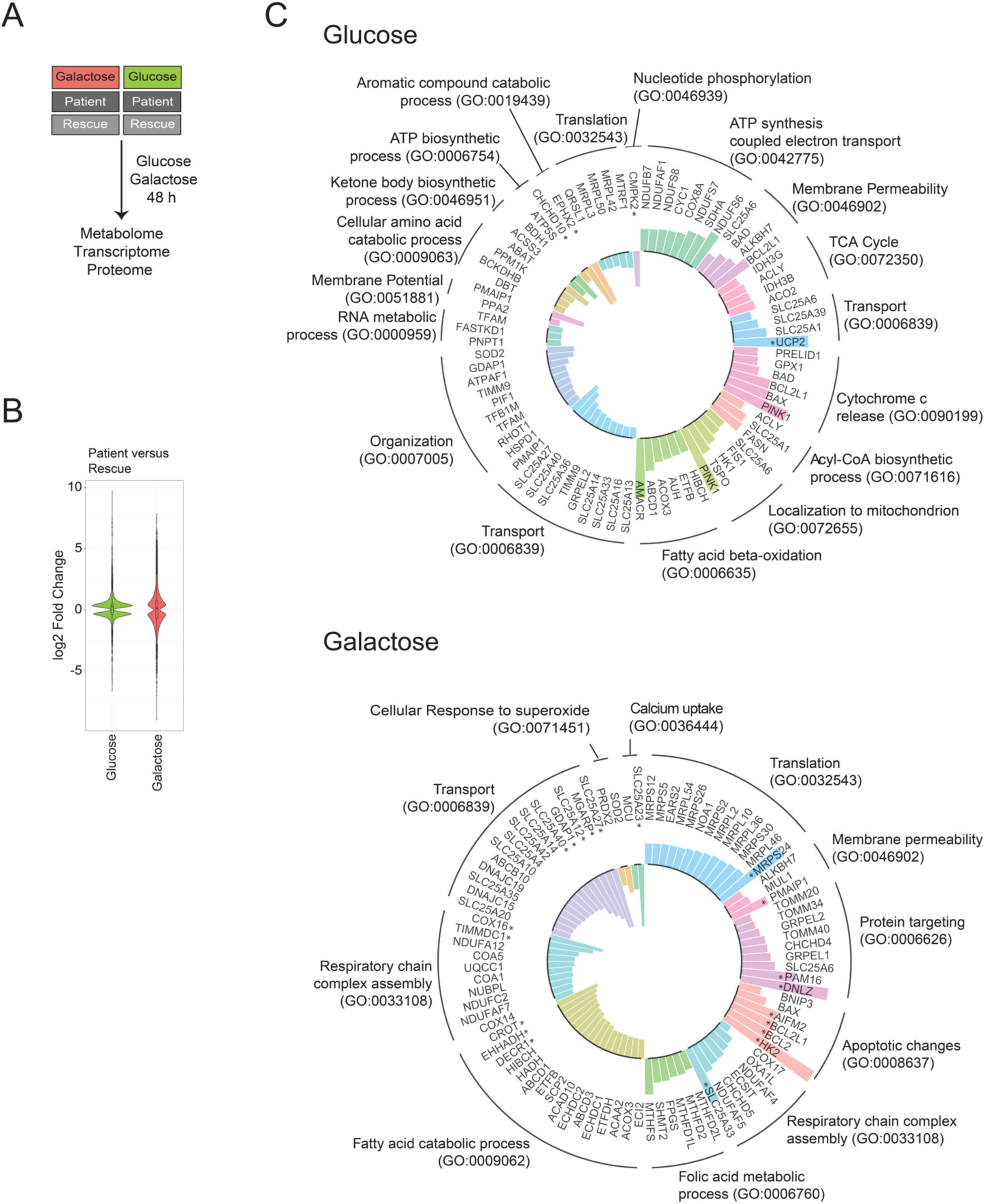
Altered biological processes revealed by RNAseq analysis. (A) Schematic representation of the omics experiments. Patient and rescue cells were grown in glucose or galactose media for 48 h and subsequently, the metabolome, transcriptome and proteome were analysed. (B) Violin Plot representing the distribution of log2 fold change of expression (patient vs rescue) in the two conditions (‘Glucose’ and ‘Galactose’) for transcripts with an adjusted p-value <0.05 (FDR<0.05). (C) Circular Bar Plot depicting mitochondrial transcripts with a minimum log2 fold change (patient vs rescue) of ±0.3 in ‘Glucose’ and ‘Galactose’, grouped based on the GO term: Biological Processes. Bars facing upwards from the inner circle represent the upregulated transcripts, and bars facing inwards represent the downregulated transcripts. Asterisk ‘*’ indicates transcripts with a log2 fold change >1.

### RNAseq analysis reveals reorganization of one-carbon metabolism

To investigate potential transcriptional alterations in patient cells and the response to galactose stress we first determined steady-state mRNA levels by RNA sequencing. The differentially expressed transcripts in ‘Glucose’ and ‘Galactose’ media showed a substantially different distribution; more transcripts had a larger fold change in ‘Galactose’ compared to ‘Glucose’ (Fig. 1B, S1A and B, Table S1). We first focused on transcripts for those proteins found in MitoCarta2.0 to ascertain which mitochondrial processes were affected in the patient cells (Calvo, Clauser et al., 2016).

In ‘Glucose’, enrichment analysis of differentially expressed transcripts for the gene ontology (GO) term for biological processes (BP), identified an upregulation of pathways involving ATP synthesis, the respiratory chain and the tricarboxylic acid cycle (TCA) cycle (Fig. 1C, ‘Glucose’). Interestingly, transcripts of the acyl-CoA biosynthetic process, for example ATP citrate synthase (ACLY), fatty acid synthetase (FASN), and the citrate transporter (SLC25A1) were upregulated in glucose even though the biosynthesis of fatty acids seems to be counterproductive in a state of compromised energy supply. Beta oxidation of fatty acids, a catabolic pathway, was upregulated as well. The most upregulated transcript was uncoupling protein 2 (UCP2), a protein involved in protection against oxidative stress and regulation of reactive oxygen species (ROS). Downregulated transcripts encompassed several mitochondrial transporters including SLC25A27 (UCP4), SLC25A14 (UCP5), SLC25A33, SLC25A36 (pyrimidines), and SLC25A13 (aspartate/malate). Transcripts coding for enzymes involved in ketone body metabolism, 3-hydroxybutyrate dehydrogenase 1 (BDH1) and acyl-CoA synthetase short chain 3 (ACSS3), were downregulated. Lastly, the mitochondrial UMP-CMP kinase (CMPK2), responsible for the phosphorylation of dUMP and dCMP, was downregulated, thus effecting the local supply of deoxynucleotides as essential precursors for mitochondrial DNA replication and transcription.

Mitochondrial processes downregulated in patient cells in ‘Galactose’ included the catabolism of fatty acids, transcripts of respiratory chain complex assembly, in particular the assembly of complexes I and IV, and a number of mitochondrial transporters such as SLC25A27 (UCP4) and SLC25A14 (UCP5) (Fig. 1C, ‘Galactose’). Both have the highest level of expression in the brain, where they were suggested to uncouple oxidative phosphorylation, exerting a protective role in cells exposed to mitochondrial defects (Hoang, Smith et al., 2012, Ramsden, Ho et al., 2012). Another transporter downregulated in patient cells in galactose is the aspartate/glutamate transporter SLC25A12, which catalyses the calcium-dependent exchange of cytoplasmic glutamate with mitochondrial aspartate (Thangaratnarajah, Ruprecht et al., 2014).

Upregulated transcripts in patient in ‘Galactose’ included those involved in mitochondrial translation, protein targeting, apoptosis, and the assembly of the respiratory chain complexes. However, the transcripts with a log2 fold change >1 mainly involved apoptotic pathways (Fig. 1C, ‘Galactose’). We also observed an upregulation of transcripts of the mitochondrial folate cycle, linked to cellular one-carbon metabolism, and which is in part responsible for nucleic acid production and therefore cellular growth. A more detailed analysis of gene transcripts involved in one-carbon metabolism revealed several changes in this pathway in the patient (Fig. 2A, B). Transcripts for mitochondrial proteins involved in folate synthesis, MTHFD2L, MTHFD2, MTHFD1L and SHMT2 were all upregulated, whereas transcripts for the cytosolic part of the one-carbon cycle, DHFR, SHMT1, TYMS and MTHFD1 were downregulated in galactose. Mitochondrial folate metabolism is responsible to supply the cytosol with formate for purine synthesis, and for the synthesis of NADPH in mitochondria (Martinez-Reyes & Chandel, 2014, Zheng, Lin et al., 2018). The same tendency was observed comparing patient cells in glucose versus patient cells in galactose, indicating a marked shift in one carbon metabolism in patient cells in galactose in particular (Fig. S1C, D). Lastly, transcripts of the serine metabolism were downregulated in the ‘Glucose’ and ‘Galactose’ conditions, indicating that serine is not synthesized *de novo* from glucose but rather directly obtained from the culture media or through the conversion of glycine to serine (Fig. 2A, B).

**Figure 2.**
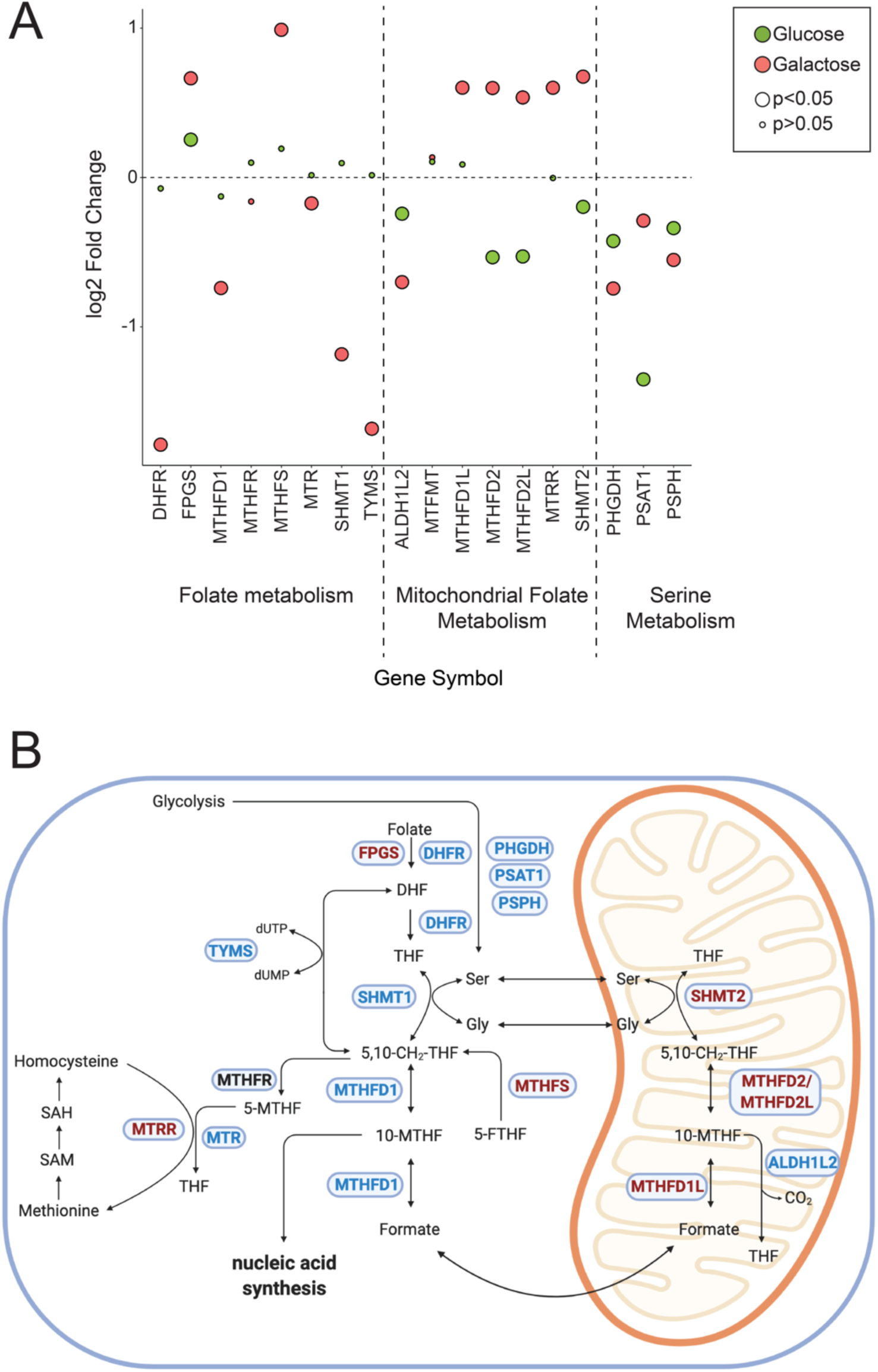
RNAseq analysis reveals changes in one-carbon metabolism. (A) Dot plot representing the fold change of the transcripts of the folate and serine metabolism. Fold change is represented in log2 scale. Large circles represent significant values (FDR<0.05), small circles show non-significant values (FDR>0.05). ‘Glucose’ (green) represents the comparison of patient versus rescue cells in glucose. ‘Galactose’ (red) represents the comparison of patient versus rescue cells in galactose. (B) Graphical representation of enzymes and metabolites of the one-carbon metabolism. Selected enzymes are shown in blue boxes for cytosolic and mitochondrial folate metabolism as well as serine and methionine metabolism. Upregulated transcripts in patient cells in galactose are shown in red and downregulated transcripts in blue. This scheme was created with BioRender.

In summary, in ‘Glucose’ patient cells upregulated transcripts of the mitochondrial respiratory chain, the TCA cycle and fatty acid beta-oxidation to compensate for the lack of mitochondrial energy production. In ‘Galactose’ on the other hand, transcripts of mitochondrial translation and mitochondrial protein import were upregulated to rescue the energy deficit. Transcripts of the one-carbon metabolism in mitochondria were upregulated to increase the supply of formate from mitochondria to the cytosol for the production of purines and to supply mitochondria with more NADPH. Interestingly, the mitochondrial aspartate/glutamate carrier was downregulated, which could lead to a deficiency of energy equivalents in mitochondria and a lack of mitochondrial aspartate in the cytosol.

### Metabolomic analysis reveals a high NADH/NAD^+^ ratio which stalls the TCA cycle in patient cells

Metabolomic analyses were performed in the four groups described above on whole cells. Using a targeted assay for 116 metabolites involved in bioenergetics, oxidative stress and mitochondrial function, the levels of 85 metabolites were successfully determined across all samples. Multi-variant data analysis using orthogonal transformation through principal component analysis (PCA) or hierarchical clustering with heatmap analysis showed tight clustering of the replicates and clear differences among the four conditions analysed, demonstrating the quality of the data (Fig. 3A and S2A). The first component PC1, showed that the treatment effect of glucose medium versus galactose medium was clearly separated, particularly for the patient cells, explaining 55% of the variance in the dataset. Moreover, we were able to separate the effect of the genotype within both the galactose or glucose condition, through the second component analysis PC2, explaining 18% of the variance. The genotype difference was larger in ‘Galactose’ than in ‘Glucose’ (Fig. 3A). The fold-change distribution of differentially detected metabolites in the two conditions ‘glucose’ and ‘galactose’ was larger in the galactose condition showing log2 fold changes ranging from +3 to -2.5 in galactose versus +1 to -2.5 in glucose (Fig. 3B, Table S2).

**Figure 3.**
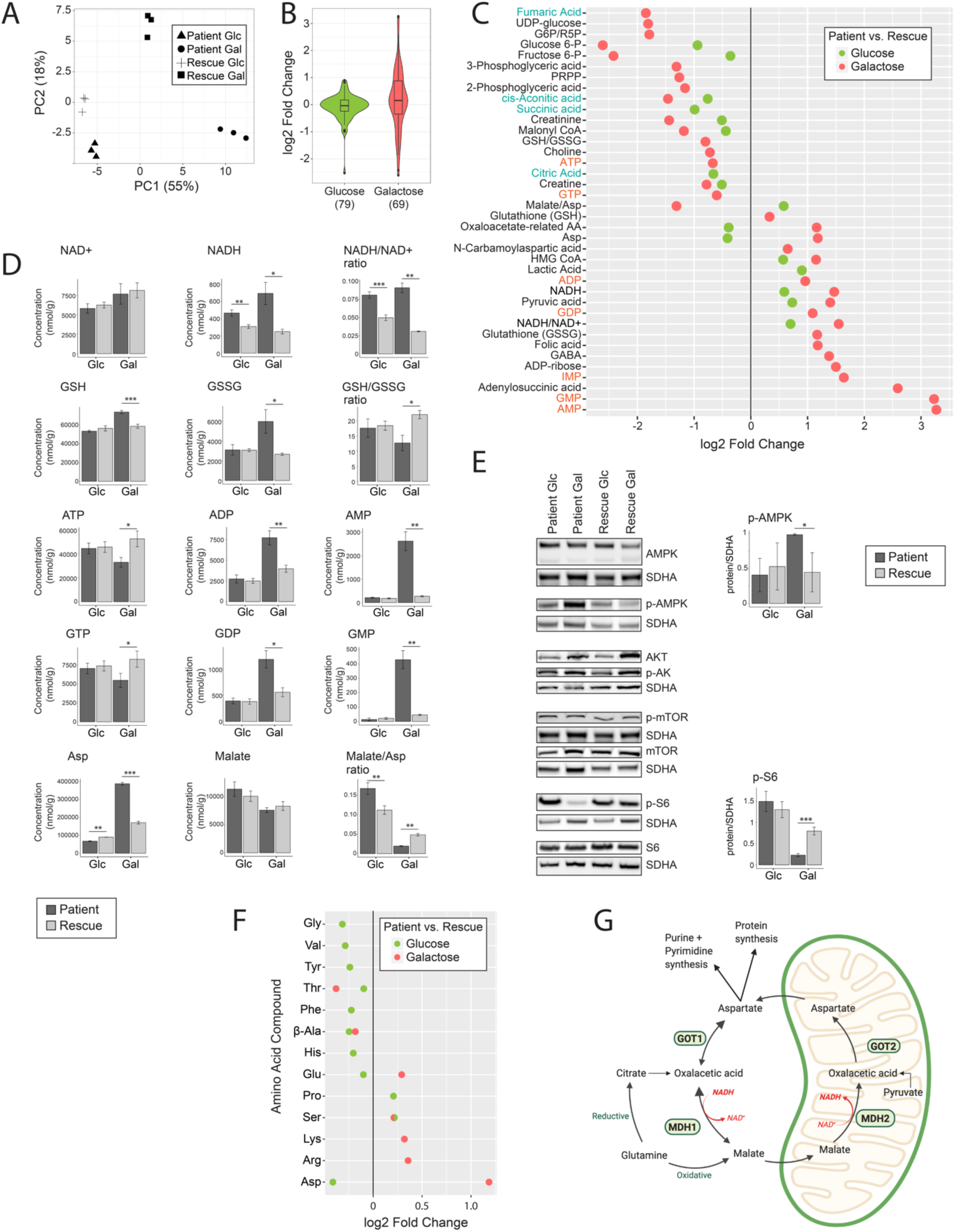
Metabolomic analysis reveals a high NADH/NAD^+^ ratio which stalls the TCA cycle in patient cells. (A) Principal component analysis of the overall metabolomic profile. Symbols convey information about the four individual groups and their three replicates. (B) Violin Plot representing the distribution of the log2 fold change (patient/rescue) for all measured metabolites. Box plots indicate the median and the first and third quartile. The number of metabolites in each group are indicated in parentheses below the graph. (C) Dot plot representing the fold change of the most changed metabolites for each condition. Fold change is represented in log2 scale and the data show only metabolites with significant differences with a p-value <0.05. The p-value is computed by Welch’s test. Metabolites of nucleotides are highlighted in orange font and TCA cycle intermediates in blue font. (D) Bar plots representing the most significantly changed metabolites. Concentrations are depicted in nmol/g and ratios are pure ratios. Values for the patient cells are in dark grey and values for the rescue cells are in light grey. P-values were computed by Welch’s test, *p<0.05, **p<0.01, ***p<0.001. (E) Immunoblot analysis of proteins involved in the mTOR pathway. Whole cell extracts from patient and rescue fibroblasts grown in glucose and galactose for 2 days were separated by SDS-PAGE and probed with antibodies against indicated proteins. SDHA was used as a loading control. Bar plot indicates the quantification of three and four immunoblot analyses for p-AMPK and p-S6, respectively, where patient is depicted in dark grey and rescue in light grey, *p<0.05, **p<0.01, ***p<0.001. (F) Dot plot representing the fold change of the most changed amino acid levels for each condition. Fold change is represented in log2 scale and data show only metabolites with significant differences with a p-value <0.05. The p-value is computed by Welch’s test. (G) Graphical representation of enzymes and metabolites of the malate aspartate shuttle. Selected enzymes are shown in green boxes for cytosolic and mitochondrial processes. This scheme was created with BioRender.

#### Redox imbalance caused by high levels of NADH in the patient

The metabolomic analysis revealed that the NADH/NAD^+^ ratio was significantly increased in both ‘Glucose’ and ‘Galactose’ in patient cells. Normally, when control cells are put in galactose the NADH/NAD^+^ ratio decreases (Ryall, Dell’Orso et al., 2015), and this was evident in the rescued cell line (Fig. 3C, D). The complex I deficiency in the patient cells likely accounted for the increase in the NADH/NAD^+^ ratio in galactose.

Additionally, the glutathione antioxidant defence pathway was activated in patient cells in galactose. Total levels of glutathione were increased (GSH + GSSG), and the GSH:GSSG ratio was decreased (Fig. 3C, D), potentially to deal with increased oxidative stress in galactose.

#### The TCA cycle is stalled in the patient

Analysis of the common upregulated and downregulated metabolites in glucose and galactose, revealed decreased levels of TCA-cycle intermediates, including cis-aconitic acid, succinate, fumarate, and citrate (Fig. 3C, Table S2). However, the level of pyruvate, a substrate entering the TCA cycle was increased (Fig. 3C). The TCA cycle is regulated by ATP and NADH levels (Berg JM, 2002), which were decreased and increased, respectively (Fig. 3C). The conversion of pyruvate into acetyl-CoA, acetyl-CoA into citrate and ketoglutarate to succinate is NADH-dependent and likely inhibited in the patient, as NADH levels were high. ADP activates the conversion from pyruvate into acetyl-CoA, acetyl-CoA into citrate and isocitrate into ketoglutarate, and its levels were increased, producing counteracting signals (Fig. 3C, D). Overall, however, the data suggested that flux through the TCA cycle was decreased in both ‘Glucose’ and ‘Galactose’.

#### High levels of aspartate do not rescue the stalled TCA cycle

With the exception of aspartate, other detected amino acids were not significantly altered in the patient cells in either ‘Glucose’ or ‘Galactose’ (Fig. 3F). Aspartate plays a key role in mitochondrial metabolism and is involved in the shuttling of electrons across the inner mitochondrial membrane through the malate-aspartate shuttle (Fig. 3G) (Berg JM, 2002). The malate/aspartate ratio thus serves as an indirect indicator of the NADH/NAD^+^ ratio and the energy state of the cell. A dysregulation of the malate/aspartate ratio was observed in both glucose and galactose; however, in opposite directions (Fig. 3D, F). The shuttle is responsible for producing NADH in mitochondria where it is reoxidized to NAD^+^ by respiratory chain complex I. The mitochondrial asparate/glutamate transporter SLC25A12 was downregulated in patient cells in galactose at the transcript level (Fig. 1C), which could lead to diminished transport of aspartate outside of mitochondria and a reduction of NADH inside of mitochondria. In the cytosol, aspartate is converted to malate by GOT1 and MDH1 (Birsoy, Wang et al., 2015). Interestingly, the enzyme GOT1 was increased at the transcript level in the patient in galactose. GOT1 normally metabolizes aspartate to oxalacetic acid, which is then converted by MDH1 into malate. This cycle allows the supply of electrons for the mitochondrial respiratory chain. When the electron transport chain is inhibited however, GOT1 produces aspartate, which serves as a substrate to synthesize purines and pyrimidines, compensating for the lack of mitochondrial aspartate synthesis (Birsoy et al., 2015, Lane & Fan, 2015) (Fig. 3G). The increased levels of aspartate and GOT1 therefore suggest that aspartate is produced by GOT1 in the cytosol to support the proliferation of patient cells. At the same time, the mitochondrial pyrimidine carrier SLC25A33 is markedly upregulated in patient cells in galactose (Fig. 1C), suggesting an increase for pyrimidine import into mitochondria.

#### Reduced energy equivalents in the patient in galactose

The metabolomic analysis revealed that nucleotide mono- and diphosphates (IMP, AMP/ADP and GMP/GDP) were the most highly increased metabolites in patient cells in ‘Galactose’, resulting in a marked increase in the AMP/ATP and GMP/GTP ratios (Fig. 3C, D), and reflecting impared mitochondrial ATP production due the complex I deficiency. Concomitantly, the metabolite adenylosuccinic acid, required for the conversion of IMP to AMP, was also markedly increased as well as ADP-ribose and folate, which are involved in the biosynthesis of purines and pyrimidines (Fig. 3C, D). Formyl folate is the precursor for inosine monophosphate, which itself is the precursor of GMP and AMP. In addition, the level of S-adenosylmethionine (SAM) a metabolite of the methionine cycle, which is linked to the folate cycle, was increased (Fig. S2B). The level of its counterpart S-adenosylhomocysteine (SAH) was below the detection limit (Fig. S2B). We conclude that increased folate levels lead to an increased generation of nucleotides IMP, AMP, and GMP, as major enzymes of the folate cycle in the cytosol were transcriptionally downregulated (Fig. 2A, B).

#### The energy balance is tightly monitored in patient cells in galactose

Concomitant with the markedly disturbed AMP/ATP ratio, we detected an increase in Thr172-phosphorylated AMP-activated protein kinase (pAMPK), signaling low cellular energy levels (Herzig & Shaw, 2018) (Fig. 3E). Another sensor of cellular energy balance is the mechanistic/mammalian target of rapamycin (mTOR) pathway. mTOR1 is a serine/threonine kinase that responds to a number of physiological inputs including nutrients (amino acids), oxygen level, hormones and energy status (AMP/ATP ratio) (Bond, 2016). Generally, mTORC1 is activated when the energy supply is sufficient, and is negatively regulated by pAMPK. We observed the inhibition of mTORC1 pathway through reduced phosphorylation of ribosomal protein S6 in patient cells in ‘Galactose’ (Fig. 3E). The levels of AKT (serine/threonine kinase), phosphorylated AKT, mTOR and phosphorylated mTOR were unchanged (Fig. 3E).

In summary, NADH/NAD^+^ ratios were high in both ‘Glucose’ and ‘Galactose’, which translated into a marked cellular energy deficit in ‘Galactose’. Glycolysis rescued the inefficient energy supply from mitochondria in ‘Glucose’; however, in ‘Galactose’ glycolysis was undetectable as expected. High levels of NADH stalled the TCA cycle as the amounts of NADH produced could not be used by the respiratory chain to generate ATP. Subsequently, entry substrates of the TCA cycle, like pyruvate and aspartate accumulated. Aspartate was additionally generated by GOT1 in the cytosol possibly in an attempt to rescue the proliferation defect of the patient in galactose.

### Integration of proteomics and metabolomics confirms reduction in TCA cycle activity at the protein level

Changes in the proteome of isolated crude mitochondria from patient and rescue cells were determined by quantitative proteomic analysis using TMT-technology. We identified 1901 unique proteins in all conditions, of which 414 proteins were mitochondrial based on MitoCarta2.0 (Calvo et al., 2016), accounting for 21.8% of the total proteins in the data set (Fig. 4A). Applying a significance level of 5% (false discovery rate, FDR), we identified differentially expressed proteins for each condition (Fig. 4B, S3, Table S3). In ‘Glucose”, several complex I subunits were downregulated in glucose (NDUFV1, NDUFS8, NDUFA7, NDUFB8) confirming the results from our first study (Straub et al., 2018) (Fig. S3). Among the most significantly upregulated proteins were four proteins in the glycolytic pathway, glyceraldehyde-3-phosphate dehydrogenase (GAPDHS), fructose-bisphosphate aldolase A (ALDOA), phosphopyruvate hydratase (ENO1), and phosphoglycerate kinase 1 (PGK1) supporting the increased activity of the glycolytic pathway in patient cells in glucose. Phosphoserine aminotransferase 1 (PSAT1), responsible for the conversion of the glycolytic intermediate 3-phosphoglycerate, an intermediate of glycolysis, to serine (Fig. 2B), was also upregulated in ‘Glucose” (Fig. S3). Serine is used through the folate cycle to produce purines. Lastly, glutathione S-transferase P (GSTP1) and glutathione S-transferase omega 1 (GSTO1), two detoxification enzymes that catalyse the conjugation of glutathione to chemical mutagens and protect against products of oxidative stress, were upregulated (Fig. S3).

**Figure 4.**
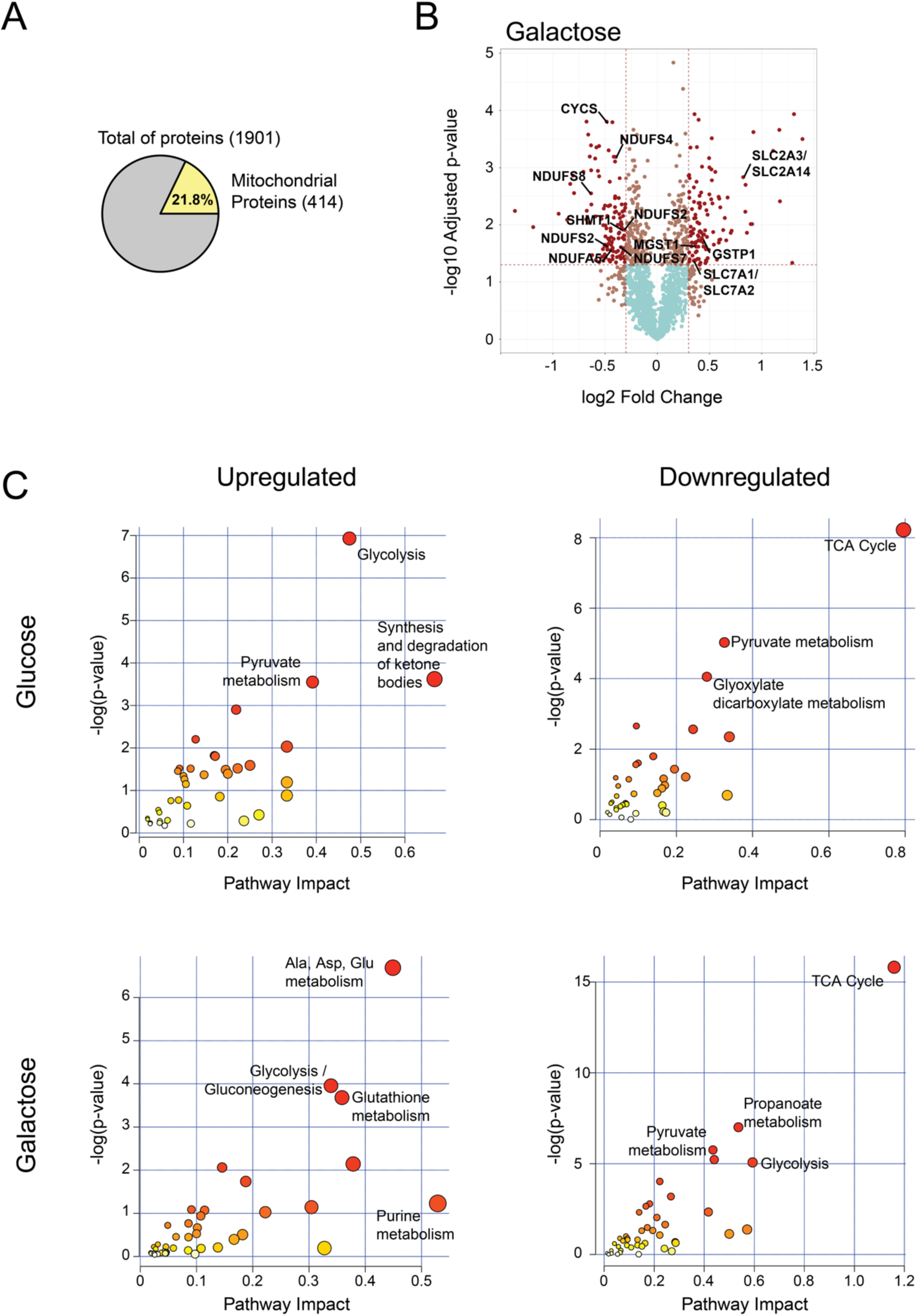
Integration of proteomics and metabolomics confirms stalled TCA cycle at the protein level. (A) Pie chart representing the total number of proteins identified in all conditions by TMT quantitative proteomics analysis. Number of nuclear encoded mitochondrial proteins (MitoCarta2.0) are indicated, as well as the percentage relative to the total. (B) Volcano plot representing the results of differential protein expression analysis between patient and rescue cells in galactose. x-axis values represent the log2 fold change and y-axis values represent the -log10 of the adjusted p-value. Selected mitochondrial proteins are labelled. (C) Joint pathway enrichment analysis performed with the MetaboAnalyst tool of significant up- or downregulated metabolites and proteins in glucose and galactose condition. Scatterplots represent p-values from integrated enrichment analysis and impact values from pathway topology analysis using KEGG metabolic pathways. The node colour represents the p-values and the node radius is based on the pathway impact values calculated with degree centrality.

In ‘Galactose’ we observed the downregulation of several complex I subunits (NDUFS8, NDUFS7, NDUFB10, NDUFA5, NDUFS5, NDUFA10, NDUFS4, NDUFS2) as well as cytochrome c (CYCS) (Fig. 4B). Interestingly, many lysosomal proteins were downregulated (ASAH1, HEXB, CTSZ, CLTB, GNS, LAMP1, CTSL, NPC2, CTSK, PSAP, PPT1, DNASE2, CTSD, LGMN, CTSB, ATP6V1G1), in particular cathepsins with cysteine-type peptidase activity (Table S3). The cytosolic serine hydroxy methyltransferase (SHMT1), which is responsible for converting serine to glycine and vice-versa, providing one-carbon units for the synthesis of methionine, thymidylate, and purines, was downregulated in ‘Galactose’ at the protein level as well at the transcript level (Fig. 4B and 2A, B).

The most prominent group of proteins upregulated in ‘Galactose’ are implicated in ER stress (Table S3). Furthermore, several transporters responsible for providing glycolysis and the TCA cycle with substrates, SLC2A1/A3 and SLC2A14 (glucose carrier) and SLC7A1/A2 (arginine, lysine and ornithine transporter) were upregulated. Lastly, we observed an increase in proteins of glutathione metabolism, GSTP1, microsomal glutathione S-transferase (MGST1), and thioredoxin domain containing 12 (TXNDC12) (Fig. 3B, Table S3).

To improve the depth of the proteomic study, we used the ‘MetaboAnalyst’ tool for the integration of proteomic and metabolomic data, which allows one to perform pathway enrichment analysis based on differentially detected metabolites and expressed proteins (Fig. 4C). We used the degree of the node (number of nodal connections) to define our enrichment analysis. Nodes with more connections, so-called hubs, have a higher impact in the pathway and, when affected, influence the enrichment more than a less connected node. Glycolysis (ALDOA, LDHA, ENO1, GAPDHS, PGK1, lactic acid, pyruvic acid), the synthesis and degradation of ketone bodies (ACAT2, 3-hydroxy-3-methylglutaryl-CoA), and pyruvate metabolism (LDHA, ACAT2, lactic acid, pyruvic acid) were the most highly induced pathways in ‘Glucose’ (Fig. 4C, S3). The increase of 3-hydroxy-3-methyl-glutaryl-CoA (HMG-CoA) (Fig. 3C) indicates a stalled mevalonate pathway, responsible for the synthesis of cholesterol and other isoprenoids. The rate limiting step of the mevalonate pathway is in the conversion of HMG-CoA to mevalonate by HMG reductase, which itself is inactivated by high levels of AMP. Therefore, in addition to TCA cycle stalling due to high levels of NADH, there was a reduction of anabolic cellular metabolism due to high levels of AMP. The integration of proteins with metabolites revealed a downregulation of the TCA cycle (DLD, OGDH, MDH2, cis-aconitic acid, succinic acid, citric acid), pyruvate metabolism (PKLR, MDH2, ALDH3A2, DLD, malonyl-CoA), and glyoxylate and dicarboxylate metabolism (MDH2, CAT, citric acid, cis-aconitic acid, glycine).

In ‘Galactose’ on the other hand we detected an up-regulation of alanine, aspartate and glutamate metabolism (ADSL, aspartic acid, adenylsuccinic acid, argininosuccinic acid, pyruvic acid, ureidosuccinic acid, gamma-aminobutyric acid), glutathione metabolism (ANPEP, TXNDC12, GSTP1, MGST1, oxidized glutathione, NADP) as well as glycolysis and gluconeogenesis (LDHAL6B, LDHA, ENO2, ENO3, PGAM1, PGAM4, TPI1, pyruvate) However, the absence of cellular lactate and intermediates of glycolysis at the metabolite level in ‘Galactose’ (Fig. 3C, Table S2) demonstrates that glycolysis does not occur in patient cells in these conditions. Similarly to ‘Glucose’, the TCA cycle is downregulated in ‘Galactose’ as well (DLD, SUCLA2, SUCLG1, IDH3A, MDH2, PDHA1, PDHB, FH, fumaric acid, cis-aconitic acid). The TCA cycle and mitochondrial activity are normally upregulated in galactose in control cells (Aguer, Gambarotta et al., 2011, Lane, Fu et al., 2015, Ostergaard, Olsson et al., 2001).

In summary, the TCA cycle and the mitochondrial respiratory chain complex I were downregulated in patient cells in both ‘Glucose’ and ‘Galactose’. This energy deficit could be compensated by high glycolytic activity in patient in ‘Glucose’, but not in ‘Galactose. Furthermore, high levels of AMP lead to a stalled mevalonate pathway, which is responsible for the synthesis of sterols, heme A and ubiquinone. Lastly, glutathione related proteins were upregulated in both conditions, indicating an increased need to deal with mitochondrial oxidative stress.

### Metabolic dysfunction leads to canonical ER UPR in patient cells in galactose

The upregulation of proteins involved in ER stress (Table S3), and the fact that mitochondrial and ER functions are tightly coupled, led us to analyse the unfolded protein (UPR) an integrated stress response (ISR) in both organelles. A coordinated activation expression of the mitochondrial unfolded protein response and the integrated stress response of the ER has been proposed previously (Melber & Haynes, 2018). In response to unfolded proteins in the ER, but also other cellular stress signals, three different arms of the UPR response can be activated through PERK, IRE1 or ATF6. In all cases the folding chaperone called binding immunoglobulin protein (BiP/GRP78/HSPA5) dissociates from PERK, IRE1 or ATF6, respectively, and leads to their oligomerization. The oligomerization of PERK as well as the activation of other kinases through other stress signals such as amino acid stress (GCN2), hypoxia (HRI), and double-stranded RNA through viral infection (PKR) initiates the ISR through the phosphorylation of eukaryotic translation factor 2 alpha (eIF2alpha) (Pakos-Zebrucka, Koryga et al., 2016). This results in the global attenuation of protein synthesis, and the selective translation of stress response proteins. Although transcript levels of PKR (*EIF2AK2*), and PERK (*EIF2AK3*) were upregulated, as were the levels of the PERK and GCN2 proteins in patient cells in ‘Galactose’ (Fig. 5A, B, S4B), we did not observe a change in phosphorylated PERK, GCN2 or eIF2alpha (Fig. 5B, S4B, C). The IRE1 pathway on the other hand is independent of eIF2alpha activation. Autophosphorylation of the kinase domains of IRE1 lead to stimulation of its endoribonuclease function, resulting in splicing of the mRNA of X-Box Binding Protein 1 (XBP1), removing a premature stop codon, and converting it into a transcxriptional activator (Adams, Kopp et al., 2019, Huang, Xing et al., 2019). Our data show, that the ER unfolded protein response in patient cells in ‘Galactose’ was activated through IRE1 and XBP1, with no evidence of activation of the ISR through the phosphorylation of eIF2alpha, the level of which was in fact reduced in patient cells in ‘Galactose’ (Fig. 5A, B). Protein phosphatase GADD34 (PPP1R15A), which recruits the protein phosphatase PP1 to dephosphorylate eIF2alpha and attenuate the ISR, was upregulated at the transcript level, possibly indicating a transcriptional response for prolonged UPR signalling (Fig. 5A, B). Consistent with the reduction in eIF2alpha phosphorylation, we observed an upregulation of transcripts involved in both cytoplasmic and mitochondrial translation (Fig. 1C, 5C). Persistent activation of IRE1 can result in the the phosphorylation of JNK, which promotes apoptosis (Almanza, Carlesso et al., 2019); however, the phosphorylation of both JNK isoforms p54 and p46 was completely abolished in patient cells in ‘Galactose’ (Fig. 5B), and significantly, the transcripts of several dual specific phosphatases (DUSPs) responsible for JNK dephosphorylation were upregulated (Fig. 5D). The activation of IRE1 was associated with the increased translation of several ER quality control genes including transcription factors ATF3 and ATF4, proteases, CHOP/DDIT3 and the cellular oxygen sensor EGLN3 (Fig. 5A, B, D).

**Figure 5.**
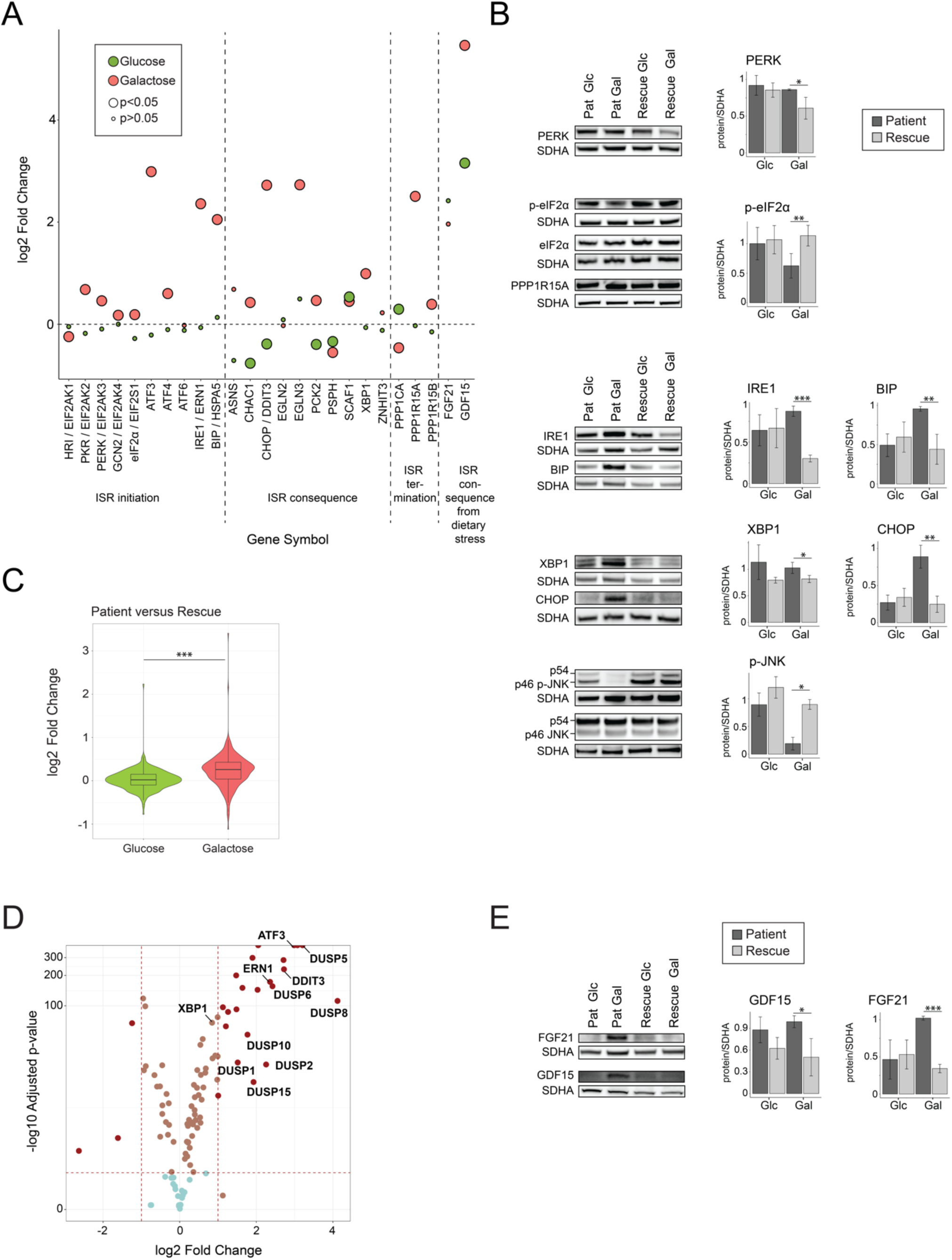
Metabolic dysfunction leads to a canonical ER stress response in patient cells in galactose. (A) Dot plot representing the fold change of the transcripts of ISR in the ER. Fold change is represented in log2 scale. Large circles represent significant values (FDR<0.05), small circles show non-significant values (FDR>0.05). ‘Glucose’ (green) represents the comparison of patient versus rescue cells in glucose. ‘Galactose’ (red) represents the comparison of patient versus rescue cells in galactose. (B) Immunoblot analysis of the proteins involved in the integrated stress response of the ER. Whole cell extracts from patient and rescue fibroblasts grown in glucose or galactose for 2 days were separated by SDS-PAGE and probed with antibodies against indicated proteins. SDHA was used as a loading control. Bar plot indicates the quantification of four (PERK), 7 (p-eIF2alpha), and three (IRE1, BIP, XBP1, CHOP, p-JNK) immunoblot analyses, normalized to SDHA, where patient is depicted in dark grey and rescue in light grey, *p<0.05, **p<0.01, ***p<0.001. (C) Violin Plot representing the distribution of the log2 fold change (patient/rescue) for transcripts involved in translation for the glucose and galactose condition. Box plots indicate the median and the first and third quartile. Wilcoxon statistical test, ***p<0.001. (D) Volcano plot representing the results of differential gene expression analysis between patient and rescue cells in galactose for proteins involved in the ISR in the ER. x-axis values represent the log2 fold change and y-axis values represent the -log10 of the adjusted p-value. Selected genes of interest are labelled. (E) Immunoblot analysis of GDF15 and FGF21. Whole cell extracts from patient and rescue fibroblasts grown in glucose or galactose for 2 days were separated by SDS-PAGE and probed with antibodies against GDF15 and FGF21. SDHA was used as a loading control. Bar plot indicates the quantification of 4 (FGF21) and 3 (GDF15) immunoblot analyses, normalized to SDHA, where patient is depicted in dark grey and rescue in light grey, *p<0.05, **p<0.01, ***p<0.001.

Two metabolic cytokines, fibroblast growth factor 21 (FGF21) and growth differentiation factor 15 (GDF15), associated with mitochondrial disease and proposed as potential biomarkers, are often activated concomitantly with stress responses (Lehtonen, Forsstrom et al., 2016, Scholle, Lehmann et al., 2018). Protein levels of both FGF21 and GDF15 as well as transcript levels of GDF15 were upregulated (Fig. 5A, E). GDF15 transcript levels were upregulated even in ‘Glucose’ (Fig. 5A); however, the upregulation of all the components of the UPR was markedly enhanced in ‘Galactose’ by comparison (Fig. 5A, S4A).

### Metabolic dysfunction leads to an activation of the mitochondrial unfolded protein response

The upregulation of the UPR in the ER as well as the increase of FGF21 and GDF15 led us to investigate the mitochondrial UPR in more detail (Fig. 6A). SIRT3, a mitochondrial NAD^+^-dependent protein deacetylase that affects gene expression in the nucleus (Scher, Vaquero et al., 2007), interacts with FOXO3 to activate anti-oxidant genes like superoxide dismutase 2 (SOD2) and catalase (CAT) (Tseng, Shieh et al., 2013). Although transcript levels of CAT and SOD2 were reduced in the patient (Fig. 6A), the level of CAT protein was increased (Fig. 6B). NAD^+^-dependent protein deacetylase SIRT7, which increases nuclear respiratory factor 1 (NRF1) activity, and alleviates mitochondrial stress by reducing the expression of mitochondrial translation proteins (Mohrin, Shin et al., 2015), was increased in our model. ATF4 and ATF5 are transcription factors that activate the mitochondrial UPR (Teske, Fusakio et al., 2013), and both were increased in patient cells in ‘Galactose’ (Fig. 6A). We also saw a marked increase in the transcript level for PPARGC1A, the transcription coactivator peroxisome proliferation-activated receptor gamma, coactivator 1 alpha (also known as PGC-1α), which enhances oxidative phosphorylation, mitochondrial biogenesis, antioxidant enzyme expression, and mitochondrial fatty acid oxidation (Fig. 6A) (Austin & St-Pierre, 2012, Cheng, Ku et al., 2018), although components of fatty acid catabolic pathway were transcriptionally downregulated (Fig. 1C). Downstream targets of the mitochondrial UPR, including proteases CLPP, HTRA2 and LONP1 and the heat shock proteins HSP70 (*HSPA9*), HSP60 (*HSPD1*) and HSP10 (*HSPE1*) were upregulated at the transcript and/or the protein level. Here too, the changes were greater in ‘Galactose’ versus ‘Glucose’ (Fig. S5). In order to clear the cells of malfunctioning organelles, autophagy is generally upregulated during the cellular stress response. In our model we could identify an upregulation of autophagy markers p62 (*SQSTM1*) and LC3 (*MAP1LC3A/B*) at the transcript and the protein level for the patient in ‘Galactose’ (Fig. 6C). This type of increase, particularly in the levels of LC3B is characteristic of a higher autophagic flux, which potentially clears damaged organelles in the patient cells (Glick, Barth et al., 2010). Lastly, we identified and increase in apoptotic cleavage of PARP and caspase 3 in patient cells in ‘Galactose’ (Fig. 6D).

**Figure 6.**
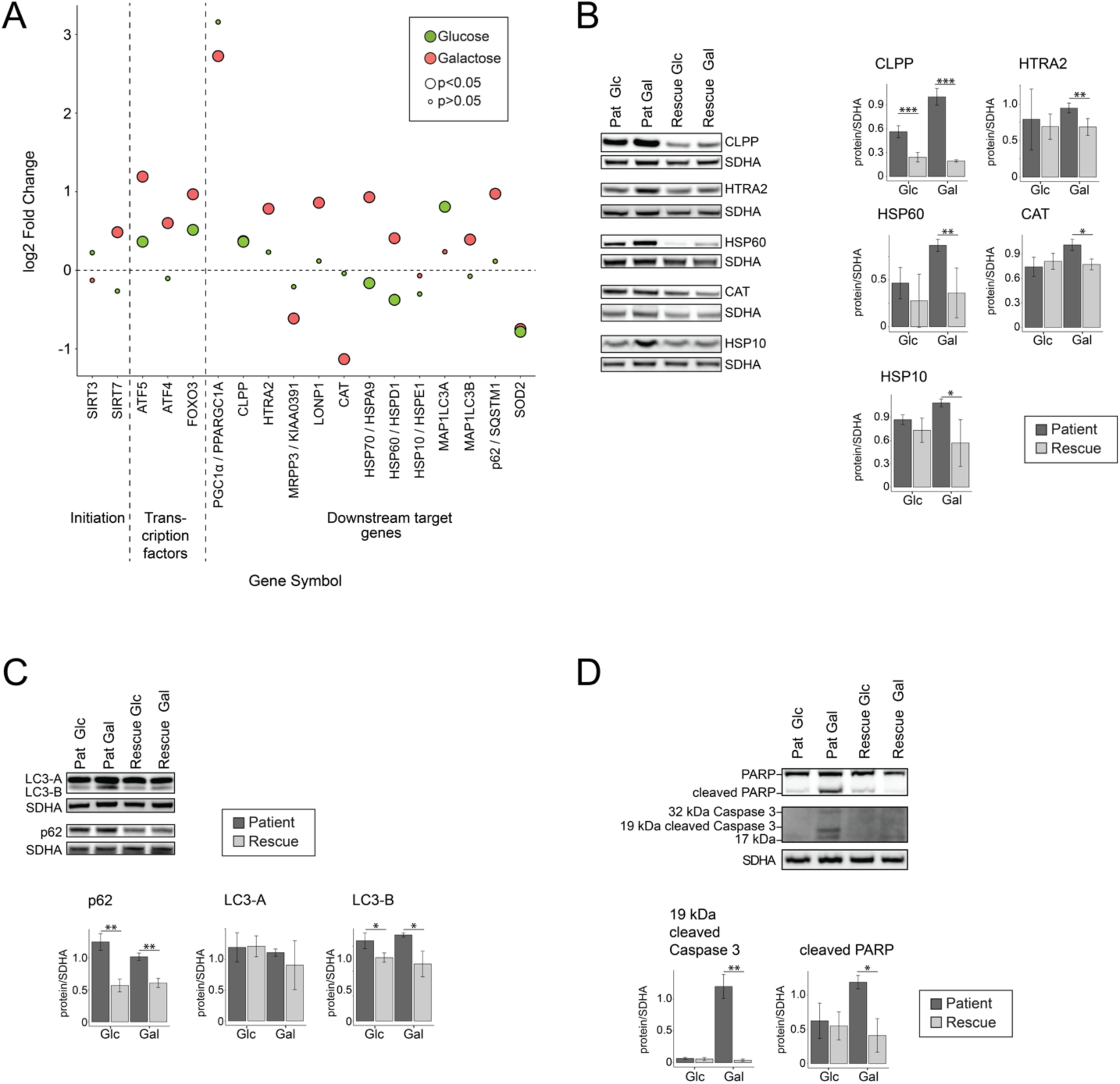
Metabolic dysfunction leads to an activation of the mitochondrial unfolded protein response (mtUPR) and increased autophagy. (A) Dot plot representing the fold change of the transcripts of mitochondrial integrated protein response. Fold change is represented in log2 scale. Large circles represent significant values (FDR<0.05), small circles show non-significant values (FDR>0.05). ‘Glucose’ (green) represents the comparison of patient versus rescue cells in glucose. ‘Galactose’ (red) represents the comparison of patient versus rescue cells in galactose. (B) Immunoblot analysis of the proteins involved in mitochondrial integrated stress response. Whole cell extracts from patient and rescue fibroblasts grown in glucose or galactose for 2 days were separated by SDS-PAGE and probed with antibodies against indicated proteins. SDHA was used as a loading control. Bar plot indicates the quantification of three (CLPP, HSP10, CAT), 4 (HTRA2, HSP60) immunoblot analyses, normalized to SDHA, where patient is depicted in dark grey and rescue in light grey, *p<0.05, **p<0.01, ***p<0.001. (C) Immunoblot analysis of the proteins involved in autophagy. Whole cell extracts from patient and rescue fibroblasts grown in glucose or galactose for 2 days were separated by SDS-PAGE and probed with antibodies against indicated proteins. SDHA was used as a loading control. Bar plot indicates the quantification of three (LC3-A, LC3-B, p62) immunoblot analyses, normalized to SDHA, where patient is depicted in dark grey and rescue in light grey, *p<0.05, **p<0.01, ***p<0.001. (D) Immunoblot analysis of the proteins involved in apoptosis. Whole cell extracts from patient and rescue fibroblasts grown in glucose or galactose for 2 days were separated by SDS-PAGE and probed with antibodies against indicated proteins. SDHA was used as a loading control. Bar plot indicates the quantification of three (cleaved PARP) and four (cleaved CC3) immunoblot analyses, normalized to SDHA, where patient is depicted in dark grey and rescue in light grey, *p<0.05, **p<0.01, ***p<0.001.

In summary, we identified the activation of the ER UPR through the IRE1/XBP1 axis, but not the ISR that is mediated through phosphorylation of eIF2alpha in patient cells in ‘Galactose’. We identified an upregulation at the transcript and protein level of specific downstream targets including CHOP, CLPP, HTRA2, HSP60, HSP10, FGF21 and GDF15 acting to rescue the patient cells in galactose, and finally Lastly, increased autophagic and apoptotic flux possibly induced by CHOP.

## Discussion

This study illustrates the metabolic consequences of nutrient stress in cells harbouring the CHCHD10 p.R15L pathogenic variant. This variant causes a fundamental metabolic reorganization with a nutrient-dependent energy deficit and an activation of the UPR in both the ER and mitochondria. The UPR response in the ER is activated through IRE1/XBP1 pathway and the mitochondrial UPR through upregulation of the transcription factors ATF4 and ATF5, both presumably acting as pro-survival mechanisms. Significantly we found no evidence for activation of an ISR, as phosphorylation of eIF2alpha was in fact decreased in the patient cells under nutrient stress. In addition, JNK appeared completely unphosphorylated, obviating pro-apototic pathways.

Our findings implicate an essential role of CHCHD10 in mitochondrial energy metabolism, directly affecting the mitochondrial respiratory chain, and subsequently resulting in a global rewiring of cellular energy metabolism. To date, six families carrying the CHCHD10 p.R15L variant (caused by c.44C>A mutation) have been reported, and all patients present with muscle wasting and progressive motor neuron disease, characteristic signs of pure ALS (Johnson et al., 2014, Khan et al., 2017, Muller et al., 2014, Zhang et al., 2015). Other CHCHD10 variants, such as p.S59L and p.G66V, cause a much broader range of disease phenotypes including frontotemporal dementia, mitochondrial myopathy and spinal muscular atrophy (Bannwarth et al., 2014, Chaussenot et al., 2014, Penttila et al., 2015, Rubino, Brusa et al., 2017). In all reported variants but one (CHCHD10 p.Q108P), disease onset is late and slowly progressive (Lehmer et al., 2018). CHCHD10 variants present with unique features of mitochondrial disease, which are rarely linked to neurodegenerative diseases like ALS and PD (Gorman, Chinnery et al., 2016).

Previous work exploring the role of CHCHD10 showed that the CHCHD10 p.R15L patient fibroblasts were unable to rely on mitochondrial respiration for energy production due to a complex I defect and therefore showed an increased use of glycolysis (Straub et al., 2018). Mitochondrial respiration in these cells could be rescued by expression of the wild-type CHCHD10 protein, suggesting that the mutation is haploinsufficient. In this study, we used the same cellular model to investigate changes in metabolites, transcripts and proteins in patient and rescued cells grown in glucose- and galactose-containing medium.

Despite numerous functional studies, the physiological role of CHCHD10 remains unknown. The deletion of the yeast orthologue *Mix17* in *S. cerevisiae* results in the reduction of mitochondrial oxygen consumption (Longen, Bien et al., 2009), implicating it in some aspect of oxidative metabolism. A link between energy metabolism and ALS has long been established, and reduced oxidative phosphorylation has been described previously for other genetic models (Jung, Higgins et al., 2002, Mattiazzi, D’Aurelio et al., 2002, Menzies, Cookson et al., 2002, Wiedemann, Manfredi et al., 2002). In human cells, siRNA mediated knockdown of CHCHD10 does not result in a destabilization of OXPHOS complexes (Burstein, Valsecchi et al., 2018); however, two CHCHD10 variants have been reported to alter the OXPHOS complex assembly/stability. The CHCHD10 p.S59L variant was shown to destabilize complex V and lead to its disassembly (Bannwarth et al., 2014), and we previously showed the complex I defect in p.R15L patient cells was phenocopied by the CRISPR-Cas9 mediated knockout of CHCHD10 (Straub et al., 2018). Here, our proteomics data confirmed the downregulation of complex I subunits in patient cells in both ‘Glucose’ and ‘Galactose’ (Fig. 4B, S3). Moreover, the transcripts for several factors involved in the complex I and IV assembly were downregulated in patient cells in ‘Galactose’ (Fig. 1C).

Nicotinamide adenine dinucleotide (NAD^+^) is an essential redox molecule, which enables mitochondrial respiration through the transport of electrons. The ratio between the oxidized and reduced form (NADH) influences many cellular and mitochondrial processes (Stein & Imai, 2012). Generally, balanced levels of NAD^+^ protect and sustain basal metabolic function and health in neurons. The depletion of NAD^+^ leads to the activation of poly (ADP-ribose) polymerases, which synthesize ADP-ribose. In our model, levels of the reducing agent NADH were upregulated in the patient, whereas NAD^+^ levels were unchanged, resulting in an increased NADH/NAD^+^ ratio. The NADH level and therefore the level of reducing equivalents in mitochondria is balanced by the malate/aspartate shuttle. In patient cells in ‘Galactose’, we identified an upregulation of the metabolite aspartate, GOT1 transcript, and a downregulation at the transcript level of the mitochondrial aspartate/glutamate carrier SLC25A12, all related to the malate-aspartate shuttle (Fig. 3G). In a state of electron transport chain deficiency and compromised aspartate export from mitochondria, which was present in patient cells in ‘Galactose’, aspartate is synthesized by reductive glutamine metabolism from glutamine, which depends on GOT1 (Metallo, Gameiro et al., 2011, Mullen, Wheaton et al., 2011) (Fig. 3G). The synthesis of aspartate, usually achieved through oxidative glutamine metabolism, is necessary for proliferation (Birsoy et al., 2015). We previously analysed the growth of patient fibroblasts 20% of which, die when the medium is switched from ‘Glucose’ to ‘Gatlactose’, while the remaining cells only start to proliferate after an adaption phase (Straub et al., 2018). This phenotype is not uncommon for cells with compromised oxidative metabolism, and has been described to be more prevalent for complex I deficiencies (Robinson, Petrova-Benedict et al., 1992, Soustek, Balsa et al., 2018). The upregulation of GOT1 in the patient cells likely indicates increased aspartate synthesis as an adaptation to the nutrient stress and to allow them to eventually proliferate. Also, an increase in mitochondrial folate metabolism could result from an increased need of formate in the cytosol, supplied by mitochondria for purine synthesis for growth (Zheng et al., 2018).

In a recent study, levels of NADH have been tied to one-carbon metabolism and cellular respiration (Yang, Garcia Canaveras et al., 2020). The TCA cycle as well as the one-carbon metabolism produce NADH in mitochondria; however, only the TCA cycle is regulated by substrate inhibition through NADH. The study showed that the accumulation of NADH due to the on-going production through the enzymatic reactions of MTHFD2/MTHFD2L in mitochondria, is toxic to cells with an impaired electron transport chain. In our model we observed an increase in transcripts of enzymes of the one-carbon metabolism and an increase in total NADH. We can therefore speculate that the regulation of NADH could present a potential target for rescuing the phenotype of CHCHD10 related disorders.

As mentioned above, the change in the NADH/NAD^+^ ratio leads to an increase in ADP-ribose, which connects the NAD pool with the AMP pool (Dolle, Rack et al., 2013). Patient cells showed changes in nucleotide levels and an energy deficit in ‘Galactose’. ATP and GTP levels were decreased whereas monophosphates IMP, AMP and GMP were markedly increased. The inability of patient cells to generate sufficient energy under the nutrient stress conditions, hints at the severity of the mitochondrial dysfunction in this model. Moreover, the imbalanced AMP/ATP ratio resulted in an increase of phosphorylated AMPK, which positively regulates signalling pathways to replenish ATP supplies (Herzig & Shaw, 2018). Mitochondrial biogenesis, autophagy and folate metabolism were upregulated to counteract the ATP deficit in patient cells in galactose. Interestingly, AMPK was previously shown to be activated in TDP-43 mouse spinal cords, and reducing AMPK activity lead to an increase in survival of motor neurons (Lim, Selak et al., 2012). Therefore, it could be advantageous to inhibit AMPK activation in order to rescue the CHCHD10 p.R15L variant induced phenotypes.

The NADH/NAD^+^ and the AMP/ATP ratios also regulate the TCA cycle (Berg JM, 2002). Depletion of ATP generally activates the TCA cycle, whereas increased levels of NADH result in reduced flux through the cycle. We identified a downregulation of metabolites and enzymes of the TCA cycle in patient cells in both ‘Glucose’ and ‘Galactose’, with more enzymes being affected in ‘Galactose’. At the same time, potential entry substrates of the TCA cycle, aspartate and pyruvate accumulated, whereas levels of entry substrates of glycolysis, glucose-6-phosphate and fructose-6-phosphate were largely decreased in patient in ‘Galactose’. The reintroduction of wild-type CHCHD10 in patient cells rescued the very low entry substrates, glucose-6-phosphate and fructose-6-phosphate; however, glycolytic activity was undetectable. Taken together, there was a tendency to decrease TCA cycle flux, aspartate and pyruvate being used for cell proliferation instead (Fig. 3G). Interestingly, glutamate in the culture media was not sufficient to upregulate the TCA cycle to normal levels.

Recent studies of two ALS mouse models have demonstrated fundamental shifts in energy metabolism. A study describing an SOD1 mouse model identified a metabolic switch from glycolysis toward the use of lipids in muscle (Palamiuc, Schlagowski et al., 2015). This switch is regulated by the pyruvate dehydrogenase kinase 4 (PDK4), which phosphorylates the PDH complex and thereby redirects the metabolic flux through the TCA cycle from lipids instead of glucose (Palamiuc et al., 2015). Another study in a mouse model of CHCHD10 p.S59L, found that white fat tissue was almost entirely lost, coinciding with an increase of the lipid metabolism regulating protein FGF21 (Anderson et al., 2019). In our model, we identified an upregulation of PDK4 at the transcript level as well as FGF21 at the protein level, consistent with the absence of glycolytic flux in ‘Galactose’. What role the lipid catabolism plays in our model could not be determined as acetoacetyl-CoA levels were not determined. However, the synthesis of lipids was possibly slightly increased in the rescue cells as the compound malonyl-CoA was increased in ‘Galactose’.

Cellular proliferation is tightly linked to cellular stress and regulated cell death. The activation of the UPR and ISR responses can be initiated through a number of specific sensors responding to stresses that include unfolded protein, viral infection, amino acid deprivation, and hypoxia (Costa-Mattioli & Walter, 2020) (Pakos-Zebrucka et al., 2016) . The activation of IRE1 leads to the splicing of XBP1, which renders the protein transcriptionally active and initiates the upregulation of stress target proteins. Moreover, IRE1 promotes a process called ‘regulated IRE1-dependent decay (RIDD), which degrades mRNAs of ER-targeted proteins to reduce the load of incoming proteins into the ER (Hollien & Weissman, 2006). The phosphorylation of eIF2alpha through PERK, PKR, GCN2 or PERK on the other hand results in an overall decrease of global cellular protein synthesis, but a specific increase of proteins that are needed for the stress response and survival (Pakos-Zebrucka et al., 2016). Recently, two groups reported the activation of the ISR for the CHCHD10 variant p.S59L and the double knock-out of CHCHD10 and CHCHD2 in mouse models (Anderson et al., 2019). CHCHD10 p.S59L accumulated as protein aggregates in a knock-in (KI) mouse and thereby induced the activation of mTORC1, the increase of transcription factors specific for the stress response, the secretion of metabolic cytokines FGF21 and GDF15, the upregulation of the serine and one-carbon metabolism, and the downregulation of the respiratory chain complexes (Anderson et al., 2019, Liu, Huang et al., 2020). Although the p.S59L CHCHD10 variant was proposed to act through a dominant gain of function mechanism (Anderson et al., 2019), and the double-knock-out through a complete loss of function, both models showed activation of the ISR. However, our results on the haploinsufficient p.R15L CHCHD10 variant show that nutrient stress elicits a UPR response in both the ER and mitochondria, and in fact a stimulation of global translation through increased dephosphorylation of eIF2alpha. This activation induced the transcription of characteristic ER and mitochondrial stress survival proteins. Moreover, we did not identify an activation of mTORC1, but rather an inhibition through the phosphorylation of AMPK. Furthermore, we observed increased autophagic and apoptotic flux, possibly mediated by increased levels of CHOP, but not phosphorylated JNK. Phosphorylated JNK is involved in apoptosis through the phosphorylation of BcL2 family of proteins (Tsuruta, Sunayama et al., 2004), and JNK1 and JNK2 double knock-out cells are resistant to apoptosis. Interestingly CHCHD10 mutant cells have also been found to be less sensitive to apoptotic cell death stimuli (Genin, Plutino et al., 2016, Tournier, Hess et al., 2000). The knockdown of XBP1 led to an increase in macroautophagy *in vivo* in other ALS models, and could be a valuable starting point for future research studying CHCHD10 related disorders (Hetz et al., 2009).

In summary, our study describes how the CHCHD10 p.R15L variant affects mitochondrial metabolism, especially under nutrient stress conditions (Fig. 7). The complex I deficiency leads to an increased NADH/NAD^+^ ratio, which is exacerbated in ‘Galactose”, where the cell relies solely on mitochondria for energy production. The cell responds to the proliferation defect associated with the energy deficit through the synthesis of aspartate from malate through GOT1 and the upregulation of the mitochondrial folate metabolism to supply formate to the cytosol. This process as well as the high levels of NADH result in diminished activity of the TCA cycle. High levels of AMP, and therefore the AMP/ATP ratio, result in increased phosphorylation of AMPK with the activation of catabolic processes on one hand and the deactivation of mTORC1 on the other. These metabolic changes activate the UPR in the ER through the IRE1/XBP1 axis and the UPR in mitochondria through ATF4 and ATF5. Finally, the patient cells either survive the existing nutrient stress by adapting their metabolism over time or undergo apoptosis, through a JNK independent mechanism. We speculate, that motor neurons, the cells that are specificly targetd ALS, are more susceptible to mitochondrial dysfunction and potentially suffer damage much earlier in limited nutrient conditions than do fibroblasts. It is known that ageing results in decreased neuronal uptake of glucose, therefore rendering motor neurons harbouring CHCHD10 p.R15L variant more susceptible to cell death in the ageing individual (Yin, Sancheti et al., 2016). We therefore propose the careful consideration of targeting mitochondrial metabolism and the UPR response for potential treatments in ALS models of CHCHD10.

**Figure 7.**
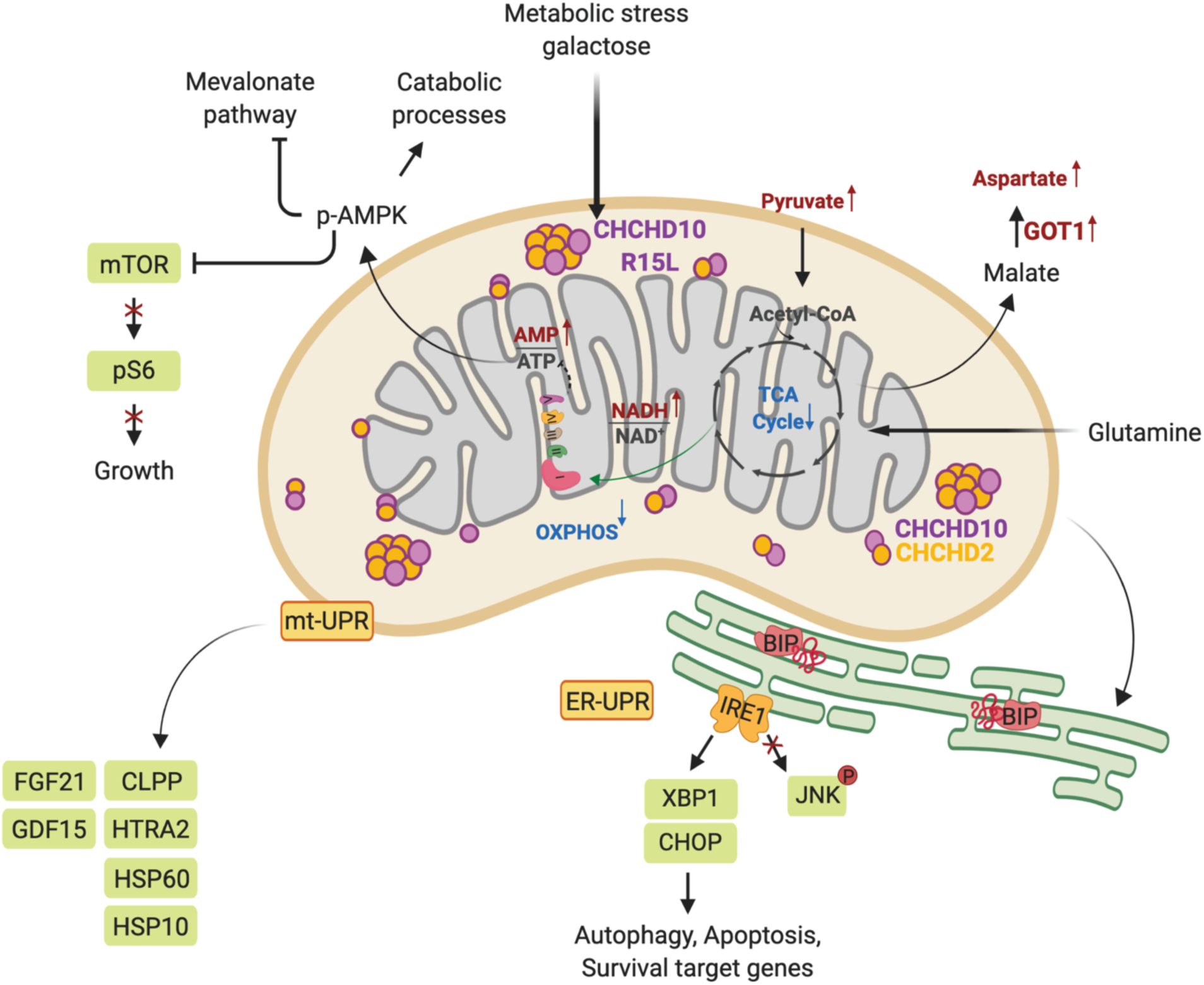
Synopsis. Mitochondrial metabolism and energy production plays a key role in cellular homeostasis. The CHCHD10 p.R15L variant leads to a complex I deficiency, resulting in an increase in the NADH/NAD^+^ ratio and a downregulation of the TCA cycle. Under nutrient stress the energy deficit in leads to an increase in the AMP/ATP ratio, increased phosphorylation of AMPK and the subsequent activation of catabolic pathways and the downregulation of the growth targeted mTOR pathway. This metabolic stress results in the activation of the UPR of the ER through IRE1/XBP1. The upregulation of different proteases and heatshock proteins, and the metabolic cytokines FGF21 and GDF15, demonstrates the simultaneous activation of the mtUPR. Finally, patient cells show increased cell death through apoptotic processes independent of JNK. This scheme was created with BioRender.

## Materials and Methods

### Human studies

The investigation of cell lines was approved by the institutional review board of the Montreal Neurological Institute, McGill University.

### Cell lines and media

A primary fibroblast culture was previously established from a heterozygous carrier of c.44G>T (p.R15L) (patient DNA #8807) diagnosed with sporadic ALS at age 54 (involving his upper limb) and died at the age of 69 (Straub et al., 2018, Zhang et al., 2015). The variant CHCHD10 p.R15L, has been associated with ALS in several independent studies (Johnson et al., 2014, Muller et al., 2014, Zhang et al., 2015). Fibroblasts stably overexpressing *CHCHD10* in patient cells (rescue) were engineered using retroviral vectors as described previously (Lochmuller, Johns et al., 1999, Straub et al., 2018). Cells were cultivated in 4.5 g/l glucose DMEM supplemented with 10% fetal bovine serum (FBS) and 5% pen/strep, at 37 °C in an atmosphere of 5% CO_2_. For carbon source-dependent experiments, fibroblasts were cultivated for two days in DMEM with 10% dialyzed FBS supplemented with either 4.5 g/l glucose or 4.5 g/l galactose before harvesting for ‘omics’ experiments.

### Mitochondrial isolation

Fibroblasts were washed twice with phosphate buffered saline (PBS), resuspended in ice-cold buffer (250 mM sucrose,10 mM Tris–HCl, pH 7.4), and homogenized by nitrogen cavitation under 500 psi pressure for 5 min. A post-nuclear supernatant was obtained by centrifugation of the samples twice for 10 min at 600 × *g*. Mitochondria were pelleted by centrifugation for 10 min at 10 000 × *g* and washed once in the same buffer. Protein concentration was determined using the Bradford assay.

### Antibodies

Antibodies directed against the following proteins were used in this study: SDHA (abcam, ab14715), AMPK (Cell Signaling, 2532), p-AMPK (Thr172) (Cell Signaling, 2535), AKT (Cell Signaling, 9272S), p-AKT (Ser473) (Cell Signaling, 4060S), mTOR (Cell Signaling, 2983P), p-mTOR (Ser2448) (Cell Signaling, 5536P), S6 (Cell Signaling, 2217), p-S6 (Ser235/236) (Cell Signaling, 4858), PERK (Cell Signaling, 5683), p-PERK (Thr980) (Cell Signaling, 3191), GCN2 (Cell Signaling, 3302), p-GCN2 (Thr899) (Abcam, ab75836), eIF2α (Cell Signaling, 2103), p-eIF2α (Ser 51) (SIGMA, SAB4504388), GADD34/PPP1R15A (Abclonal, A17117), IRE1/ERN1 (Cell Signaling, 3294), BIP (abcam, ab21685), XBP1 (Santa Cruz, sc-7160), CHOP/DDIT3 (Cell Signaling, 5554), JNK (Cell Signaling, 9252), p-JNK/SAPK (T183/Y185) (Cell Signaling, 9251), FGF21 (abcam, ab171941), GDF15 (abcam, ab106006), CLPP (Proteintech, 15698-1-1AP), HTRA2 (Proteintech, 15775-1-AP), HSP60 (Santa Cruz, sc-136291), CAT (Abclonal, A11777), HSP10 (Santa Cruz, SC-376313), LC3 (Cell Signaling, D3U4C), SQSTM1/p62 (Cell Signaling, 8025), PARP (Cell Signaling, 9542), cleaved PARP (Cell Signaling, 9541), cleaved caspase 3 (Cell Signaling, 9661S).

### SDS-PAGE and immunoblot analysis

Cells were pelleted and lysed with 1.5% *n*-dodecyl-D-maltoside (DDM) in PBS with cOmplete™ protease inhibitor (Roche) for 15 min on ice. Cells were centrifuged at 20,000 × g for 20 min at 4 °C. Protein concentration was determined using Bradford assay. 2x Laemmli buffer was added to 20-30 μg of protein and denatured at 55 °C for 15 min. The mixture was run on denaturing 8 % to 12.5 % polyacrylamide gels. Separated proteins were transferred onto a nitrocellulose membrane and immunoblot analysis was performed with the indicated antibodies in 5% milk in Tris-buffered saline and Tween 20 (TBST).

### RNA extraction and RNA sequencing

Total RNA from fibroblasts was extracted and purified by using RNeasy Mini kit (QIAGEN). RNA quality was tested on a 1% agarose gel and then RNA sequencing was performed by GENEWIZ on the Illumina HiSeq instrument.

### Transcriptomic analysis

The sample analysis was done by GENEWIZ. The original read counts were normalized to adjust for various factors such as variations of sequencing yield between samples. These normalized read counts were used to accurately determine differentially expressed genes. Data quality assessments were performed to detect any samples that are not representative of their group, and thus, may affect the quality of the analysis. Using DESeq2, a comparison of gene expression between groups of samples was performed. Groups were defined as follows: (1) ‘Glucose’: patient in glucose versus rescue in glucose, (2) ‘Galactose’: patient in galactose versus rescue in glucose, (3) ‘Patient’: patient in galactose versus patient in glucose and (4) ‘Rescue’: rescue in galactose versus rescue in glucose. The Wald test was used to generate p-values and log2 fold changes. Genes with an adjusted p-value < 0.05 and absolute log2 fold change > 1 were considered as differentially expressed genes. RNA sequencing data was deposited to NCBI Gene Expression Omnibus (GEO), GSE144725.

### Gene ontology analysis

Gene ontology (GO) analysis of significant differentially expressed transcripts was performed using the annotation tool ENRICHR (https://amp.pharm.mssm.edu/Enrichr/, GO_BP 2018), focusing on the GO term of biological processes (Chen, Tan et al., 2013, Kuleshov, Jones et al., 2016). As an input, gene symbols of genes passing the cut off of log2 fold change of ≷ ± 0.3 were used. A maximum p-value of 0.05 was chosen to select only significant enrichment.

### Metabolite extraction and analysis

Metabolite extraction was performed according to the protocol of Human Metabolome Technologies, Inc. Briefly, 5 million cells where seeded in four replicates for each condition, one for counting and three for the collection of metabolites in three independent biological replicates. The culture medium was aspirated from each culture plate and cells were washed twice with washing buffer (5% w/w mannitol solution). The washing buffer was aspirated and methanol (LC/MS grade) together with an internal standard solution was added to extract metabolites. Cell extracts were centrifuged for 5 min at 2,300 × g at 4 °C. The supernatant was transferred into ultracentrifuge filter cups and centrifuged at 9,000 × at 4 °C for 3 hours. The tube was set up in a centrifugal evaporator for 6 hours to evaporate the supernatant and obtain the metabolites as precipitate. Metabolites were quantified by Human Metabolome Technologies, Inc. Metabolic pathway analysis was performed using the integrative tool for joint pathway analysis in the MetaboAnalyst 4.0 platform (http://www.metaboanalyst.ca; (Chong, Soufan et al., 2018)). Metabolites and proteins were selected with a cut-off of log2 fold change of ≷ ± 0.2 and a p-value < 0.05. Selected parameters were: hypergeometric test for enrichment analysis, degree centrality for topology analysis, metabolic pathways for pathway database and ‘combine query’ as the integration method.

### Proteome extraction and analysis

Fibroblasts were grown to 90% confluency in 15 cm dishes to lead to a minimum of 2 × 10^6^ cells per condition. Cells were rinsed 2-3 times with 1x PBS to remove cell culture media and pelleted at 600 × g. Mitochondria were isolated as described above and lysed with lysis buffer (50 mM HEPES, 1% SDS, 150 mM NaCl). The lysate was centrifuged at 16,000 × g for 10 minutes at 4 °C. Protein concentration was determined in the supernatant by BCA Protein Assay Kit (Thermo Fisher Scientific, #23227). 100 μg of protein was transferred into a new tube and the final volume was adjusted to 100 μL with 100 mM triethylammonium bicarbonate (TEAB). 5 μL of 200 mM tris(2-carboxyethyl)phosphine) (TCEP) was added and samples were incubated at 55° C for 1 hour. 5 μL of 375 mM iodoacetamide was added to the mix and let stand for 30 minutes protected from light at room temperature. 600 μL of pre-chilled acetone (−20 °C) was added and the mix was put at -20 °C over-night to precipitate. Samples were centrifuged at 8000 × g for 10 minutes at 4 °C. The tubes were carefully inverted to decant the acetone and let dry for 30 min at room temperature. Samples were digested, labelled and analysed by mass spectrometry on an Orbitrap (Thermo Scientific) at the Institute de Recherches Cliniques (IRCM) de Montreal. The isobaric labelling of the peptides was performed using 10-plex TMT reagents (Thermo Fisher Scientific).

Differentially expressed proteins were identified by performing Welch’s t-statistic test, comparing patient cells with rescue cells in glucose and galactose. Values were normalized by subtraction of the mean from the log2 values of the comparison ratios. The mass spectrometry proteomics data have been deposited to the ProteomeXchange Consortium via the PRIDE partner repository with the dataset identifier PXD018806 and 10.6019/PXD018806.

### Bioinformatic and statistical analysis

Pathways, transcripts and proteins selected in each condition were filtered after Benjamini-Hochberg correction at an adjusted p-value < 0.05 (FDR 5%). All data analysis and plots were generated using R and RStudio and modified using Adobe Illustrator CC. For the rest of the analysis, data were expressed as mean ± SEM, and p-values were calculated using two-tailed Welch’s t-test for pairwise comparisons of metabolites and proteins and the Wald as well as the Wilcoxon test for transcripts. Statistical tests were performed using RStudio and GraphPad Prism 6.0 (*, P < 0.05; **, P < 0.01; ***, P < 0.001; n≥ 3).

## Acknowledgements

We thank Hana Antonicka, Aurel Besse-Patin, and Mari Aaltonen for fruitful discussions and critique of the manuscript and Logan Walsh for the help with the analysis of the RNAseq data. This work was supported by grants from the CIHR, Parkinson Canada, and the Muscular Dystrophy Association to EAS. IS was financially supported by fellowships from Parkinson Canada and Healthy Brain for Healthy Lifes (HBHL, McGill University).

## Author contribution

IR planned, and executed all the experiments and wrote the manuscript. WW executed several immunoblot experiments. EAS supervised the project and wrote the manuscript.

## Conflict of interest statement

The authors report no conflict of interest.

## Notes

https://www.ncbi.nlm.nih.gov/geo/query/acc.cgi?acc=GSE144725

### Competing Interest Statement

The authors have declared no competing interest.

### Summary of Updates

Only the title of Table S1 was corrected.

