## Supplementary Figures and Legends for "A CHCHD10 variant causing ALS elicits an unfolded protein response through the IRE1/XBP1 pathway"

### Supplementary Materials

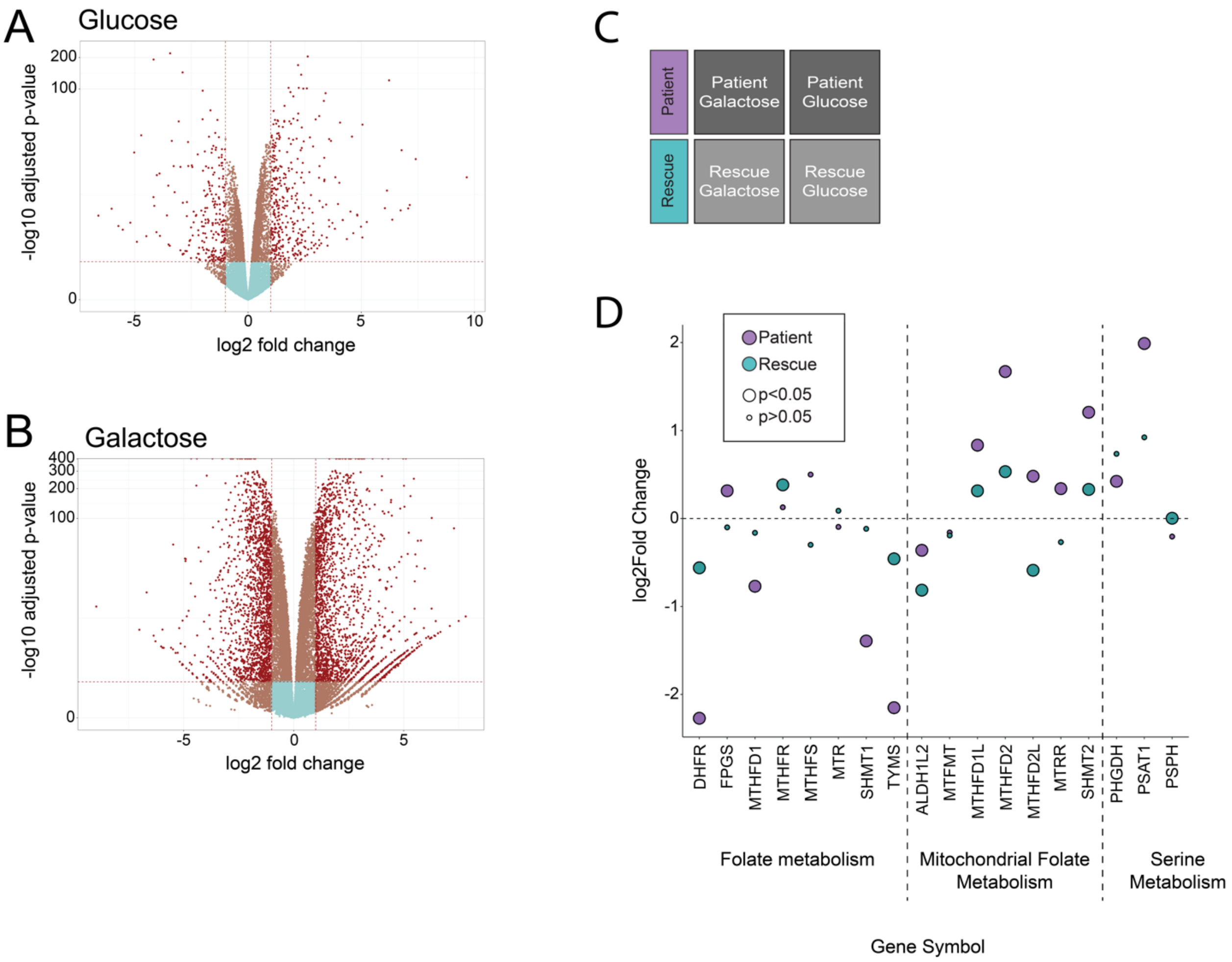


**Figure S1** Analysis of the transcriptome

(A) Volcano plot representing the results of differential transcript expression analysis between patient and rescue cells in 'Glucose'. x-axis values represent the log2 fold change and y-axis values represent the -log10 of the adjusted p-value.

(B) Volcano plot representing the results of differential transcripts expression analysis between patient and rescue cells in 'Galactose'. x-axis values represent the log2 fold change and y-axis values represent the -log10 of the adjusted p-value.

(C) Schematic representation of the treatment comparison. 'Patient' (purple) is defined as patient in galactose versus patient in glucose. 'Rescue' (blue) is defined as rescue in 'Galactose' versus 'Glucose'.

(D) Dot plot representing the fold change of the transcripts of the folate and serine metabolism in response to the treatment. Fold change is represented in log2 scale. Large circles represent significant values (FDR<0.05), small circles show non-significant values (FDR>0.05). ‘Patient’ (purple) represents the comparison of patient cells in ‘Galactose’ versus ‘Glucose”. ‘Rescue’ (blue) represents the comparison of rescue cells in ‘Galactose’ versus ‘Glucose’.

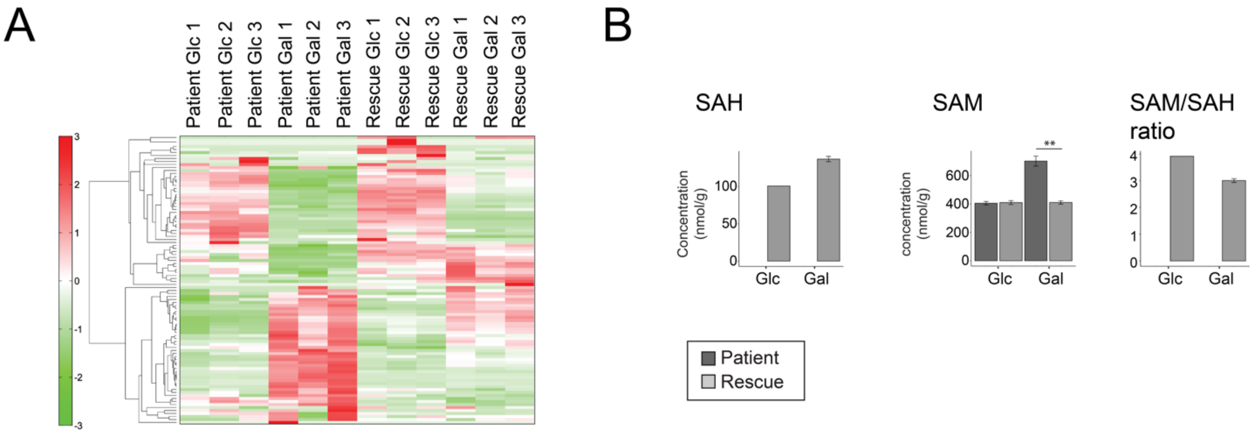


**Figure S2** Analysis of the metabolome

(A) Heat map analysis of the metabolome. Hierarchical clustering of samples (columns) and metabolites (rows) was based on the standardized value of relative area in detected peaks. Upregulated metabolites are shown in red, downregulated metabolites in green.

(B) Bar plots representing the change of SAH, SAM and their ratio. Concentrations are depicted in nmol/g and ratios are pure ratios. Values for the patient cells (dark grey), value for the rescue cells (light grey). The p-value is computed by Welch’s test. SAH was not detected in patient cells.

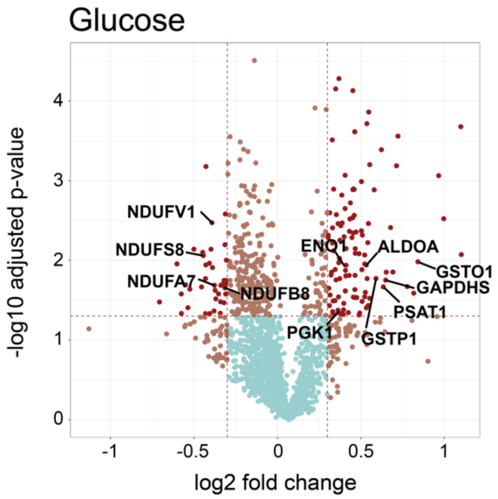


**Figure S3** Proteome in glucose

Volcano plot representing the results of differential protein expression analysis between patient and rescue cells in glucose. x-axis values represent the log2 fold change and y-axis values represent the -log10 of the adjusted p-value. Selected mitochondrial proteins of interest are labelled.

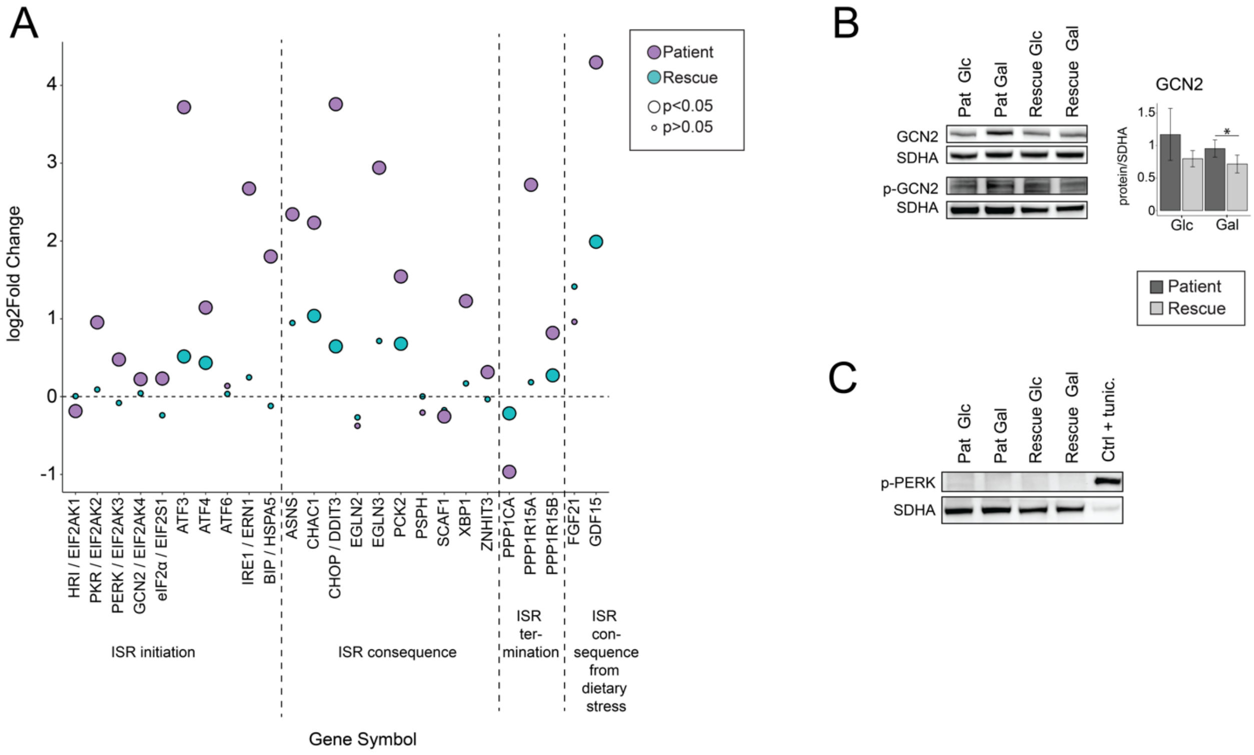


**Figure S4** Canonical ER stress response in patient and rescue cells in response to treatment

(A) Dot plot representing the fold change of the transcripts of the ISR and UPR responses in response to treatment. Fold change is represented in log2 scale. Large circles represent significant values (FDR<0.05), small circles show non-significant values (FDR>0.05). 'Patient' (purple) represents the comparison of patient cells in 'Galactose' versus "Glucose'. 'Rescue' (blue) represents the comparison of rescue cells in 'Galactose' versus 'Glucose'.

(B) Immunoblot analysis of GCN2 and phosphorylated GCN2. Whole cell extracts from patient and rescue fibroblasts grown in glucose or galactose for 2 days were separated by SDS-PAGE and probed with antibodies against indicated proteins. SDHA was used as a loading control. Bar plot indicates the quantification of 7 (GCN2) immunoblot analyses, normalized to SDHA, where patient is depicted in dark grey and rescue in light grey, *p<0.05, **p<0.01, ***p<0.001.

(C) Immunoblot analysis of phosphorylated PERK. Whole cell extracts from patient and rescue fibroblasts grown in glucose or galactose for 2 days were separated by SDS-PAGE and probed with antibodies against indicated proteins. SDHA was used as a loading control. A control sample was treated with 1 μg/mL tunicamycin for 12 hours as positive control.

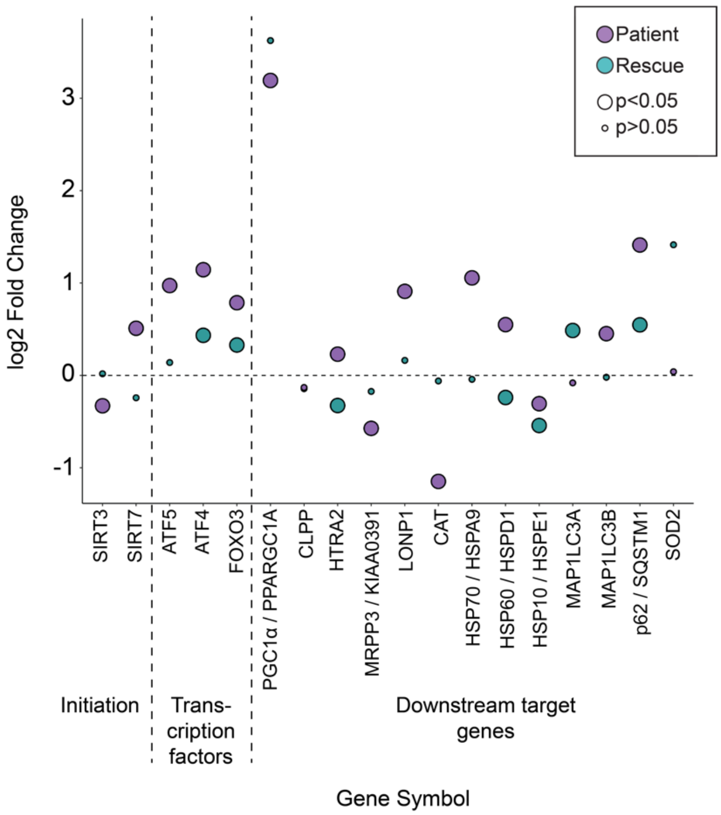


**Figure S5** Canonical mtUPR in patient and rescue cells in response to treatment

Dot plot representing the fold change of the transcripts of mitochondrial UPR in response to treatment. Fold change is represented in log2 scale. Large circles represent significant values (FDR<0.05), small circles show non-significant values (FDR>0.05). ‘Patient’ (purple) represents the comparison of patient cells in ‘Galactose’ versus ‘Glucose’. ‘Rescue’ (blue) represents the comparison of rescue cells in ‘Galactose’ versus ‘Glucose’.

**Table S1** Differential expression analysis of transcripts from RNA sequencing analysis.

**Table S2** Absolute concentrations of metabolites in patient and rescue cells grown in 'Glucose' or 'Galactose' and the subsequent metabolomic analysis.

**Table S3** Total intensities of proteins from TMT-proteomic experiment and differential expression analysis.
